# BREEDIT: A novel multiplex genome editing strategy to improve complex quantitative traits in maize (*Zea mays* L.)

**DOI:** 10.1101/2022.05.02.490346

**Authors:** Christian Damian Lorenzo, Kevin Debray, Denia Herwegh, Ward Develtere, Lennert Impens, Dries Schaumont, Wout Vandeputte, Stijn Aesaert, Griet Coussens, Yara de Boe, Kirin Demuynck, Tom Van Hautegem, Laurens Pauwels, Thomas B. Jacobs, Tom Ruttink, Hilde Nelissen, Dirk Inzé

**Affiliations:** Center for Plant Systems Biology, VIB, B-9052 Gent, Belgium; Department of Plant Biotechnology and Bioinformatics, Ghent University, B-9052 Gent, Belgium; Flanders Research Institute for Agriculture, Fisheries and food (ILVO), B-9820 Merelbeke, Belgium

**Keywords:** Multiplex gene editing, CRISPR/Cas9, multiplex amplicon sequencing, maize, gene family, network engineering, reverse genetics

## Abstract

Ensuring food security for an ever-growing global population while adapting to climate change is the main challenge for agriculture in the 21^st^ century. Though new technologies are being applied to tackle the problem, we are approaching a plateau in crop improvement using conventional breeding. Recent advances in gene engineering via the CRISPR/Cas technology pave the way to accelerate plant breeding and meet this increasing demand. Here, we present a gene discovery pipeline named ‘BREEDIT’ that combines multiplex genome editing of whole gene families with crossing schemes to improve complex traits such as yield and drought resistance. We induced gene knockouts in 48 growth-related genes using CRISPR/Cas9 and generated a collection of over 1000 gene-edited maize plants. Edited populations displayed, on average, significant increases of 5 to 10% for leaf length and up to 20% for leaf width compared with controls. For each gene family, edits in subsets of genes could be associated with increased traits, allowing us to reduce the gene space needed to focus on for trait improvement. We propose BREEDIT as a gene discovery pipeline which can be rapidly applied to generate a diverse collection of mutants to identify subsets of promising candidates that could be later incorporated in breeding programs.

## Introduction

The production of enough food to feed the increasing global population is facing many challenges due to climate change. Extreme temperature ranges, reduction of water availability and limited use of arable land are all expected to converge on a significant drop in crop yields (Zhang and Cai, 2011; Long et al., 2015; Brás et al., 2021). During the past century, conventional breeding has been decisive to adapt crops to local environments and to increase yield under stress conditions (Nuccio et al., 2018; Snowdon et al., 2021). Genomics-assisted breeding has greatly contributed to generate new varieties by incorporating haplotype information in breeding programs (Bhat et al., 2021). Nonetheless, we are slowly approaching a plateau in crop improvement using conventional breeding, since gene discovery and introgression of favorable alleles cannot be implemented fast enough to cope with the losses caused by environmental stresses.

In that perspective, innovative strategies need to be implemented to bridge the gap between conventional breeding and the knowledge acquired through plant molecular biology to further improve complex traits such as yield. Crop yield is determined by the complex interaction of the (a)biotic environment with the genetically determined growth and developmental processes that drive the plant’s life cycle (Elias et al., 2016). There are numerous yield-related traits such as early seedling vigor, root and shoot architecture, biomass allocation, resource use efficiency, senescence, seed filling, etc. In some cases, such as disease resistance, few causative genes control the expression of the trait. However, for many yield- and growth-related quantitative traits (e.g. organ growth, tolerance to abiotic stress such as drought), numerous, small-effect genes contribute (Mickelbart et al., 2015; Poland and Rutkoski, 2016). Traditionally, yield improvement has been tackled from two distinct angles. Breeding aims at producing genetic combinations with better performance, whereas molecular biology works to understand the mode of action of yield-related genes. These two fields operate at very different scales; breeding recombines chromosomal segments towards a favorable genome constitution, whereas molecular biology only deals with a limited number of genes. In crop breeding programs, phenotypes (e.g. seed yield) are collected from many individuals and multi-year/multi-location field trials. By correlating the phenotypes with the genotypic diversity of individuals, genetic variants associated with the improved trait values can be identified (Rasheed et al., 2017). Using this approach, many quantitative agronomic traits have been found to be determined by numerous small-effect loci, with the underlying genomic regions known as ‘quantitative trait loci’ or QTLs. Such QTLs are generally searched for in segregating mapping populations of recombinant inbred lines (RILs) obtained from two or more parents. A more recent variant of this approach is the genome-wide association study (GWAS), in which numerous genome-wide markers are assayed in many diverse genotypes to associate loci with the phenotypic trait (Wang and Qin, 2017). Furthermore, the combination of phenotypic trait data with the availability of a high number of genomic markers, or even the entire genome sequence, can be used for genomic prediction to increase the predictability of the breeding value of new material (Voss-Fels and Snowdon, 2016). Although these marker-assisted breeding technologies have a major impact on the accuracy and speed of crop breeding, the genes underlying the QTLs are in many cases unknown. In recent years, technological advances have combined GWAS with molecular -omics phenotypes that go beyond the genomic information, so that molecular networks start to emerge in molecular breeding (Baute et al., 2015; Baute et al., 2016; Xiao et al., 2016; Miculan et al., 2021).

Over the past four decades, there has been tremendous progress in the understanding of the molecular basis of many different plant processes. The use of model organisms such as *Arabidopsis* and rice has been a driving force. A vast amount of research delivered insights into the molecular pathways steering seed development, root growth, leaf development, plant architecture, tolerance to severe drought stress, cold tolerance, flooding and many more agronomic traits. Combined, this information reinforced the idea that plant growth and possibly crop yield may be improved by altering the expression of specific (regulatory) genes. Indeed, many reports have shown that positive effects of yield-related traits could be obtained by modifying the expression of individual genes. In *Arabidopsis*, more than 60 genes were identified that, when ectopically expressed or down-regulated, increase leaf size and in many cases also the size of other organs, including seeds (for reviews, see Gonzalez et al., 2012; Czesnick and Lenhard, 2015; Vercruysse et al., 2020). Likewise, numerous genes that can be used to improve seed yield and size in rice have been identified (Li and Li, 2016). Based on these observations, agro-biotech companies have initiated large-scale programs in the beginning of the 21^st^ century to investigate the effect of numerous selected genes on agronomic traits in crops of interest, mainly maize and rice. The conclusion of these studies was that although positive effects were often noticed in the greenhouse and even in field trials, the observed changes were often too small and too much dependent on the genotype and the environment to justify further investments in pursuing this high-throughput screening approach (Paul et al., 2018; Simmons et al., 2021). Why is it so challenging to translate basic insights in molecular networks and genes into improved crops? In breeding, the phenomenon of expressivity is well-known. Expressivity measures the extent to which a given genotype is expressed at the phenotypic level. The concept of expressivity is best explained by the notion that genes often work in complex networks with many different levels of regulation. Such higher-order regulation is typically exerted on complex and essential processes, such as growth, that need to integrate a panoply of endogenous, genetically determined signals as well as environmental cues. Single-point perturbations of networks often have a limited effect because other components of the network take over to buffer the system. However, in many cases, the combination of perturbations of a network makes phenotypes much more visible. For example, the pairwise combinations of 13 *Arabidopsis* growth-related genes (GRGs), each enhancing leaf size on their own when ectopically expressed or mutated, lead in more than 80% of the combinations to additive or synergistic effects on leaf size (Vanhaeren et al., 2014; Vanhaeren et al., 2017). Moreover, a triple combination of three different mutants of GRGs increased the size of leaves, flowers, seeds and even roots of Arabidopsis in a spectacular manner (Vanhaeren et al., 2017). Also in maize, albeit with fewer genes, pairwise combinations of specific alleles of growth-enhancing genes result in additive effects (Sun et al., 2017; Liu et al., 2021). This concept is also clearly observed during breeding, when yield traits most often are determined by many small-effect loci that need to work in concert to obtain a maximal output.

Despite the spectacular advances made by systems biology in integrating large data sets, the mechanisms behind the control of plant developmental processes are so complex that predicting which combination of genes would provide the optimal effect on yield remains virtually impossible. Understanding the mode of action might be the best way forward to estimate the combinability of genes (Vanhaeren et al., 2014; Sun et al., 2017). However, even when dealing with a relatively small number of genes, testing all possible pairwise gene combinations remains cumbersome and resource intensive. The investments become even more important when triple or higher-order gene combinations have to be tested, which is necessary to achieve stable yield increases of 10% or higher.

The clustered regularly interspaced short palindromic repeat (CRISPR) technology emerged as a powerful tool for simultaneously multiplex-targeting several GRGs, easily generating genetic variability in a broad set of targets and thus enabling a plethora of combinatorial mutations to be analyzed (Knott and Doudna, 2018; Zhang et al., 2019). Several studies showed how CRISPR could be used to reshape plant architecture and target complex traits in multiple species like tomato (Rodríguez-Leal et al., 2017; Wang et al., 2021), wheat (Li et al., 2020), rice (Meng et al., 2017) and in maize (Doll et al., 2019). As a broader application, large-scale CRISPR screens have been carried out in rice (Lu et al., 2017), cotton (Ramadan et al., 2021), maize (Liu et al., 2020; Gong et al., 2022), tomato (Jacobs et al., 2017), oilseed rape (Li et al., 2018) and soybean (Bai et al., 2020).

Here, we explored an experimental approach to bridge the gap between conventional breeding and genetic engineering of multiple genes by combining multiplex CRISPR-mediated genome editing with crossing schemes to observe favorable phenotypes. We denominated this approach BREEDIT, a contraction of breeding and gene editing, and propose this strategy as a powerful technique to engineer complex traits by knocking out a large number of key players in gene families and pathways. In just two generations, we generated a list of putative gene knockouts (KOs) required to evoke clear yield-related phenotypes in maize. BREEDIT could therefore be used to rapidly identify a subset of genes involved in the expression of a complex trait and identify targets for plant breeding programs.

## Results

### Development of a CRISPR/Cas9 multiplex genome editing pipeline in maize: general outline

The aim of this study was to develop a flexible pipeline that combines multiplex gene editing and different crossing schemes to generate plants with modified traits (Figure 1). First, 48 candidate GRGs (Table 1) were selected based on the literature or in-house knowledge, in the target species maize, complemented with other model organisms, i.e. *Arabidopsis* and rice (Figure 1A). For instance, negative growth regulators whose inactivation is likely to result in positive effects on growth are suitable GRG candidates. Guide RNAs (gRNAs) targeting these GRGs are designed and cloned into multiplex gene editing vectors (referred to as SCRIPTs), which are then used to transform *Cas9*-expressing lines (named EDITOR lines), resulting in super transformed lines that harbor both Cas9 and a SCRIPT containing 12 gRNAs (Figure 1B; Supplemental Figure S1). The BREEDIT pipeline then uses highly multiplex (HiPlex) amplicon sequencing combined with the SMAP haplotype-window bioinformatics workflow to routinely monitor gene edits at gRNA cutting sites. Amplicon sequencing at great depths enables to collect haplotype sequences and their respective frequencies (Figure 1D). Both types of information can be used to assess the effect of mutations on the encoded protein function or activity and assign a genotype to the plant for a specific locus. Per sample and per locus, the length difference between a mutated haplotype and the reference haplotype is used to classify the mutated haplotypes in two categories: haplotype_KO_ corresponds to haplotypes containing out-frame insertions or deletions (indels), leading to a gene KO and an nonfunctional protein; haplotype_REF_ includes haplotypes with only single-nucleotide polymorphims (SNPs) outside the cutting site or in-frame indels supposed to have less impact on the translated protein that may still behave as the reference protein. In CRISPR/Cas9 experiments, one plant may contain more haplotypes than its ploidy level because of mosaic tissues as a consequence of the initial (T0) or ongoing (T1, T2) Cas9 activity, thus complicating the genotyping. To interpret complex haplotype constitutions, the relative fraction of all haplotype_KO_ is summed per locus per sample. The resulting aggregation is interpreted as a gene loss-of-function (LOF) dosage, further discretized in three categories: LOF_0/2_ (none of the two chromosomes is affected by a set of haplotype_KO_), LOF_1/2_ (one of the two chromosomes is affected by a set of haplotype_KO_), and LOF_2/2_ (both chromosomes are affected by a set of haplotype_KO_). The three dosage categories are used in genotype-to-phenotype associations.

**Figure 1.**
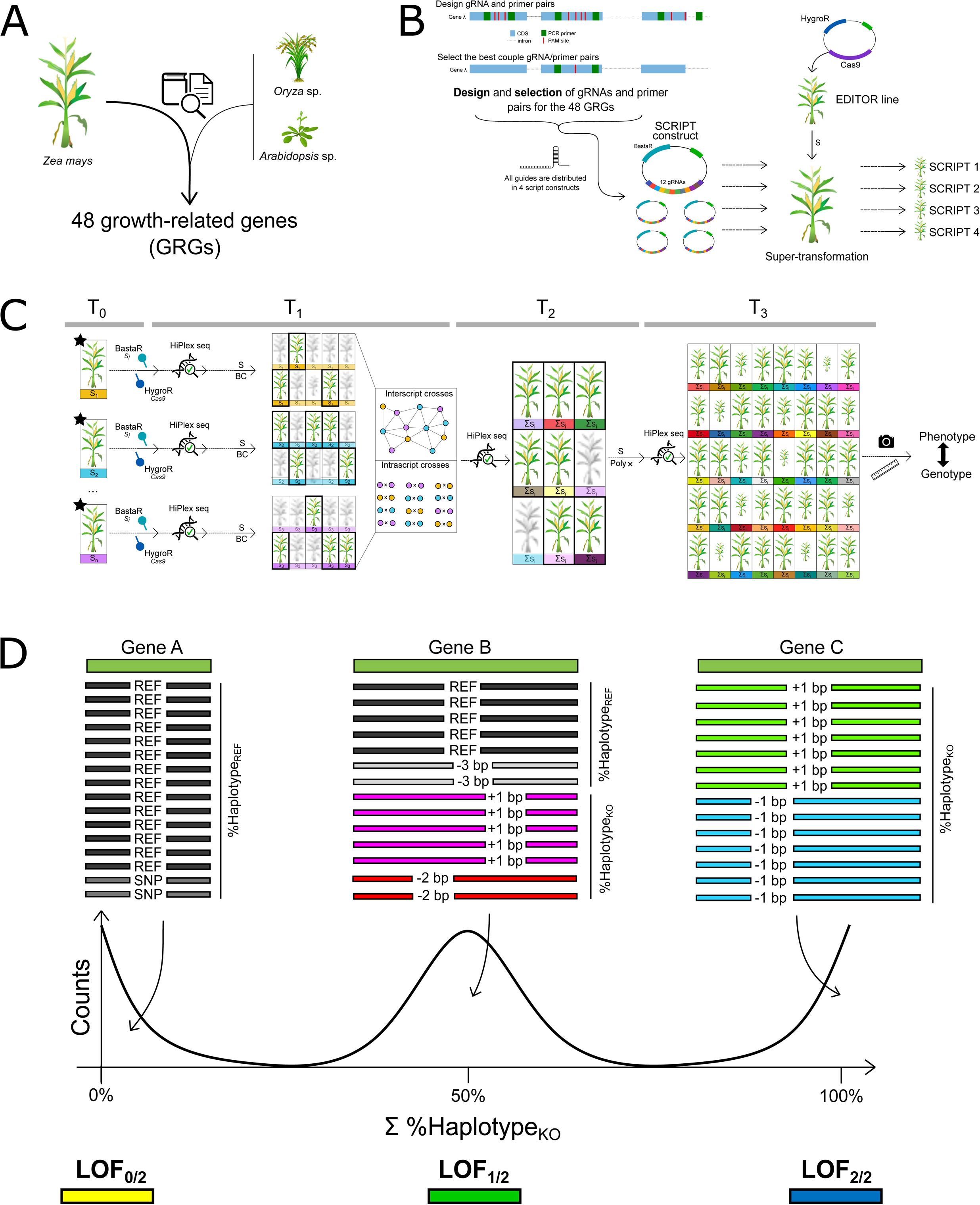
The multiplex gene editing strategy of BREEDIT. **A**. Selection of growth-related genes (GRGs) is based on published and in-house research performed in Arabidopsis, rice or maize **B**. After gene selection, gRNAs with NGG PAM sites are selected for each gene and PCR primer pairs are designed to re-sequence gRNA target sites and flanking regions via HiPlex amplicon sequencing. Per gene, the best set of gRNA and flanking primer pairs is selected. Twelve gRNAs are cloned in multiplex gene editing vectors named “SCRIPTs”. Next, the SCRIPT constructs are transformed in a Cas9-expressing maize line named EDITOR. **C**. Vigorous T0 plants containing the SCRIPT (BASTA resistant) and the Cas9 EDITOR construct (hygromycin resistant) are further genotyped using HiPlex amplicon sequencing. Based on the genotypes, plants are selected either for crossing with B104 (BC: backcross), with plants with complementary mutations caused by the same SCRIPT (intra-script crosses) or with plants containing a different SCRIPT and therefore mutations in genes from a different family or pathway (inter-script crosses). These crosses aim at maximizing the mutation landscape and diversity. Finally, self-crosses (S) of lines generate a segregating progeny for high-throughput phenotyping of selected traits, which later can be associated to (combinations of) genes. **D**. From continuous read depth to discrete loss-of-function (LOF) genotypic classes. Sequencing reads are mapped to the B104 reference loci. Two read categories are derived, namely haplotype_REF_ and haplotype_KO_. Haplotype_REF_ corresponds to the aggregated fraction of reads containing only SNPs, in-frame indels or being the reference haplotype. Haplotype_KO_ refers to the aggregated fraction of reads with out-frame indels. A tri-modal distribution is expected for haplotype_KO_, with local maxima around 0%, 50%, 100%, each corresponding to a fraction of the genome being edited at the locus. Haplotype_KO_ is therefore discretized in three gradual classes of LOF: LOF_0/2_ (the genome is not edited with out-frame indels, i.e. 0 chromosome out of 2 in a diploid organism), LOF_1/2_ (half of the genome is edited with out-frame indels, i.e. 1 chromosome out of 2 in a diploid organism), LOF_2/2_ (all the genome is edited with out-frame indels, i.e. 2 chromosomes out of 2 in a diploid organism).

**Table 1.** List of the 48 GRGs targeted by different SCRIPTs.

| SCRIPT | Position | Gene | Gene family/pathway | B73 V3 gene id | References |
| --- | --- | --- | --- | --- | --- |
| 1 | 1 | *ZmGa2ox2* | GA2-oxidases | GRMZM2G006964 | (Huang et al., 2015; Li et al., 2021) |
| 1 | 2 | *ZmGa2ox4* | GA2-oxidases | GRMZM2G153359 |  |
| 1 | 3 | *ZmGa2ox5* | GA2-oxidases | GRMZM2G176963 |  |
| 1 | 4 | *ZmGa2ox7* | GA2-oxidases | GRMZM2G427618 |  |
| 1 | 5 | *ZmGa2ox8* | GA2-oxidases | GRMZM2G155686 |  |
| 1 | 6 | *ZmGa2ox9* | GA2-oxidases | GRMZM2G152354 |  |
| 1 | 7 | *ZmGa2ox13* | GA2-oxidases | GRMZM2G031432 |  |
| 1 | 8 | *D8* | DELLA/GRAS family | GRMZM2G144744 | (Winkler and Freeling, 1994; Lawit et al., 2010) |
| 1 | 9 | *D9* | DELLA/GRAS family | GRMZM2G024973 |  |
| 1 | 10 | *ZmSLRL1-1* | DELLA/GRAS family | GRMZM5G826526 | (Ikeda et al., 2001; Itoh et al., 2005) |
| 1 | 11 | *ZmSLRL2* | DELLA/GRAS family | GRMZM5G874545 |  |
| 1 | 12 | *ZmSPY* | GA signalling | GRMZM2G357804 | (Qin et al., 2011) |
| 2 | 1 | *ZmCKX-2* | cytokinin oxidases | GRMZM2G050997 | (Ashikari et al., 2005; Bartrina et al., 2011) |
| 2 | 2 | *ZmCKX-3* | cytokinin oxidases | GRMZM2G167220 |  |
| 2 | 3 | *ZmCKX-4* | cytokinin oxidases | GRMZM5G817173 |  |
| 2 | 4 | *ZmCKX-4B* | cytokinin oxidases | GRMZM2G024476 |  |
| 2 | 5 | *ZmCKX-5* | cytokinin oxidases | GRMZM2G325612 |  |
| 2 | 6 | *ZmCKX-6* | cytokinin oxidases | GRMZM2G404443 |  |
| 2 | 7 | *ZmCKX-7* | cytokinin oxidases | GRMZM2G134634 |  |
| 2 | 8 | *ZmCKX-8* | cytokinin oxidases | GRMZM2G162048 |  |
| 2 | 9 | *ZmCKX-9* | cytokinin oxidases | GRMZM2G303707 |  |
| 2 | 10 | *ZmCKX-10* | cytokinin oxidases | GRMZM2G348452 |  |
| 2 | 11 | *ZmCKX-11* | cytokinin oxidases | GRMZM2G122340 |  |
| 2 | 12 | *ZmCKX-12* | cytokinin oxidases | GRMZM2G008792 |  |
| 3 | 1 | *ZmKRP1;1* | ICK/KRP cyclin-dependent kinase | GRMZM2G101613 | (Cheng et al., 2013; Cao et al., 2018) |
| 3 | 2 | *ZmKRP1;2* | ICK/KRP cyclin-dependent kinase | GRMZM2G084570 |  |
| 3 | 3 | *ZmKRP1;3* | ICK/KRP cyclin-dependent kinase | GRMZM2G343769 |  |
| 3 | 4 | *ZmKRP3* | ICK/KRP cyclin-dependent kinase | GRMZM2G154414 |  |
| 3 | 5 | *ZmKRP4;2A* | ICK/KRP cyclin-dependent kinase | GRMZM2G037926 |  |
| 3 | 6 | *ZmKRP4;2B* | ICK/KRP cyclin-dependent kinase | GRMZM2G116885 |  |
| 3 | 7 | *ZmKRP5;1* | ICK/KRP cyclin-dependent kinase | GRMZM2G358931 |  |
| 3 | 8 | *ZmKRP5;2* | ICK/KRP cyclin-dependent kinase | GRMZM2G157510 |  |
| 3 | 9 | *ZmPP2C-A9* | ABA signal transduction | GRMZM2G134628 | (He et al., 2019) |
| 3 | 10 | *ZmPP2C-A11* | ABA signal transduction | GRMZM2G159811 |  |
| 3 | 11 | *ZmHB124B* | Homeobox transcription factor family | GRMZM2G023291 | (McConnell et al., 2001) |
| 3 | 12 | *ZmHB124C* | Homeobox transcription factor family | GRMZM2G178102 |  |
| 4 | 1 | *ZmTCP3* | TCP - CIN clade | GRMZM2G115516 | (Koyama et al., 2017; Sarvepalli and Nath, 2018; Lan and Qin, 2020) |
| 4 | 2 | *ZmTCP8* | TCP - CIN clade | GRMZM2G020805 |  |
| 4 | 3 | *ZmTCP9* | TCP - CIN clade | GRMZM2G589470 |  |
| 4 | 4 | *ZmTCP10* | TCP - CIN clade | GRMZM2G166946 |  |
| 4 | 5 | *ZmTCP22* | TCP - CIN clade | GRMZM2G120151 |  |
| 4 | 6 | *ZmTCP25* | TCP - CIN clade | GRMZM2G035944 |  |
| 4 | 7 | *ZmTCP42* | TCP - CIN clade | GRMZM2G180568 |  |
| 4 | 8 | *ZmGRF4* | Growth-regulating factor clade | GRMZM2G004619 | (Nelissen et al., 2012; Voorend et al., 2016; Liebsch and Palatnik, 2020) |
| 4 | 9 | *ZmGRF10* | Growth-regulating factor clade | GRMZM2G096709 |  |
| 4 | 10 | *ZmGRF17* | Growth-regulating factor clade | GRMZM2G124566 |  |
| 4 | 11 | *ZmPHD8* | SET domain transcription factor | GRMZM2G409224 | - |
| 4 | 12 | *ZmBPC6* | GAGA-binding protein | GRMZM2G118690 | (Gong et al., 2018) |

After selecting transgenic lines, T0 lines are genotyped and the T0 plants with the highest numbers of gene KOs (either partial (LOF_1/2_) or complete (LOF_2/2_)) are crossed to obtain material for phenotyping (Figure 1C). Different crosses can be performed to maximize the number of edited genes as well as to fix combinations of gene edits. Self-crosses serve to fix edits in parallel to maximize phenotypic readout, while backcrosses to the original line provide heterozygous lines which can later be self-crossed and phenotyped in T2. Additional specific crosses can be performed to further enrich edit diversity. Plants harboring the same SCRIPT but containing edits at different genes from that SCRIPT can be crossed to increase the number of gene edits (up to 12) in the corresponding gene family or pathway. Such crosses will be further referred to as intra-script crosses. Furthermore, plants transformed with different SCRIPTs can be crossed to maximally combine mutants in genes covered by different families or pathways. These crosses will be further referred to as inter-script crosses. Since Cas9 remains active in all subsequent generations (Impens et al., 2022), new transgenerational edits are expected to accumulate, resulting in a large collection of higher-order mutants (up to 24 gene edits when two SCRIPTs are combined) in different segregating states (i.e. LOF_0/2_, LOF_1/2_ or LOF_2/2_).

Because several plants are generated following the BREEDIT approach, easy-to-measure quantitative traits are used to maximize the throughput of the phenotyping steps. Despite the high number of plants generated, each individual has likely a unique genotypic profile given the many combinations of indels and dosage that can happen in a set of 12 genes or more. Therefore, repetitions of same genotypic combinations cannot be used for statistics in BREEDIT. The effects of combinations of gene edits on traits are better appraised at the population level, though the specific causative gene combination cannot be deduced. The effect of a single gene on a trait can however still be evaluated considering that multiple observations of a single gene KO would conceal the putative noise brought by mutations in other genes. The framework for phenotyping experiments consists of several (minimum two) independent trials to test the performance of independent mutated populations compared with the EDITOR line. Single-gene associations to a trait are then conducted per experiment per population. The number of times a gene KO is significantly associated with a trait across different independent populations and experiments is a measure for the importance of that gene in the expression of the trait. At the end of the BREEDIT pipeline, genes can be ranked to delineate a minimal set of candidate genes with maximal effect on trait expression, thus reducing the gene space to be considered for further research.

### Applying the BREEDIT strategy

To test the BREEDIT strategy, we selected 48 maize GRGs with potential positive effects on growth when mutated, individually or in combination (Table 1). The gRNAs targeting the 48 genes were distributed over SCRIPT 1 to SCRIPT 4 and were, as much as possible, grouped per gene family. This distribution primarily aims to simultaneously knockout multiple members of the same gene family/pathway to overcome potential functional redundancy of paralogs. In addition, grouping by family aims to generate segregating mutants with a range of gene KOs, which may help to untangle complex relationships in gene regulatory networks that might be overlooked when only single or double mutants are considered. Additionally, chromosomal positions of the GRGs were taken into consideration to spread the distribution of genes belonging to a same SCRIPT over chromosomes when possible (Supplemental Figure S2). The 12 genes targeted in SCRIPT 1 are major players in gibberellin catabolism and signaling. The 12 genes targeted in SCRIPT 2 are maize orthologs of genes encoding cytokinin oxidases (*CKXs*), key regulators of cytokinin catabolism. SCRIPT 3 contains gRNAs for eight genes encoding the family of inhibitors of cyclin-dependent kinase/Kip-related proteins (*ICK/KRP*), as well as four genes expected to encode negative regulators of growth under drought conditions: two maize *PP2C* orthologs (*ZmPP2Cs*) and two *HOMEOBOX*-type genes (*HB124B and HB124C*), orthologs of the *Arabidopsis* genes *PHABULOSA* and *PHAVOLUTA* (McConnell et al., 2001). Finally, SCRIPT 4 contains gRNAs for seven orthologs of class II *CINCINNATA*-*TEOSINTE BRANCHED 1/CYCLOIDEA/PROLIFERATING CELL FACTOR (CIN-TCP)* and three members of the *GROWTH REGULATING FACTORS* (*GRF)* genes, major regulators of cell division, leaf shape and leaf size determination. Additionally, gRNAs targeting an ortholog of the GAGA-binding protein-encoding *BASIC PENTACYSTEINE 6* (*ZmBPC6*) and a gene encoding a plant homeodomain (PHD)-finger protein (*ZmPHD8*) were included in SCRIPT 4.

### Generation of highly edited maize populations for all SCRIPTs

We developed a set of three independent homozygous EDITOR lines that constitutively express the Cas9 protein in the maize inbred line B104 (Supplemental Figure S3) to execute editing at loci targeted by arrays of 12 gRNAs expressed from the SCRIPT vector. EDITOR 1 and EDITOR 3 were supertransformed with SCRIPT 1 for a preliminary evaluation of gene editing. After transformation, the EDITOR 1 and EDITOR 3 supertransformed populations showed similar editing profiles (Supplemental Figure S4). At T0, six out of the 12 targeted genes showed LOF_1/2_ or LOF_2/2_ in both EDITOR lines and the number of mutant alleles at each locus was comparable between both EDITOR backgrounds. The same gRNAs were active in both EDITOR backgrounds but four genes out of the six being commonly edited in both EDITOR backgrounds have LOF_2/2_ in EDITOR 1 whereas two in EDITOR 3 (Supplemental Figure S4). We proceeded with EDITOR 1 as the genetic background for further experiments and supertransformed this line with the remaining three scripts. Like for SCRIPT 1, we monitored gene edits in T0 plants and all subsequent generations using HiPlex amplicon sequencing. Indels in haplotype sequences ranged from -90 bp to +92 bp. Insertions of one nucleotide (+1 bp) were the most represented type of mutation, but overall, deletions were more present than insertions (Figure 2A). The largest insertions showed sequence similarity to genomic fragments located up to 1 kb upstream or downstream of the expected cutting site. At T0, we detected haplotype_KO_ in 11, 12, 8, and 12 out of the 12 target sites for SCRIPT 1, 2, 3, and 4, respectively (Figure 2B). Across all T0 SCRIPT populations, a large diversity of haplotypes (109 haplotypes with in-frame and 407 haplotypes with out-frame indels) could be identified (Supplemental Figure S5). Some haplotype_KO_ were initially not detected at T0, but appeared in T1 populations (Figure 2B) of both intra-script and inter-script crosses, revealing either ongoing gene editing in subsequent generations or overlooked edits due to mosaic tissues in T0. Overall, from T0 to T2, mutations could be found in all the 48 targeted genes, except one (*SPY* in SCRIPT 1). We focused on haplotype_KO_, and observed a diversity of haplotype_KO_ combinations per locus per sample (mono-, bi-, multi-allelic) in the T0 to T2 samples, all expected to lead to a gene LOF, either partial (LOF_1/2_) or complete (LOF_2/2_) (Figure 2C). We observed a typical tri-modal distribution for the aggregated fraction of haplotype_KO_ that could be roughly divided into three areas with higher counts, each corresponding to a discrete genotypic class (LOF_0/2_, LOF_1/2_, and LOF_2/2_; Supplemental Figure S6).

**Figure 2.**
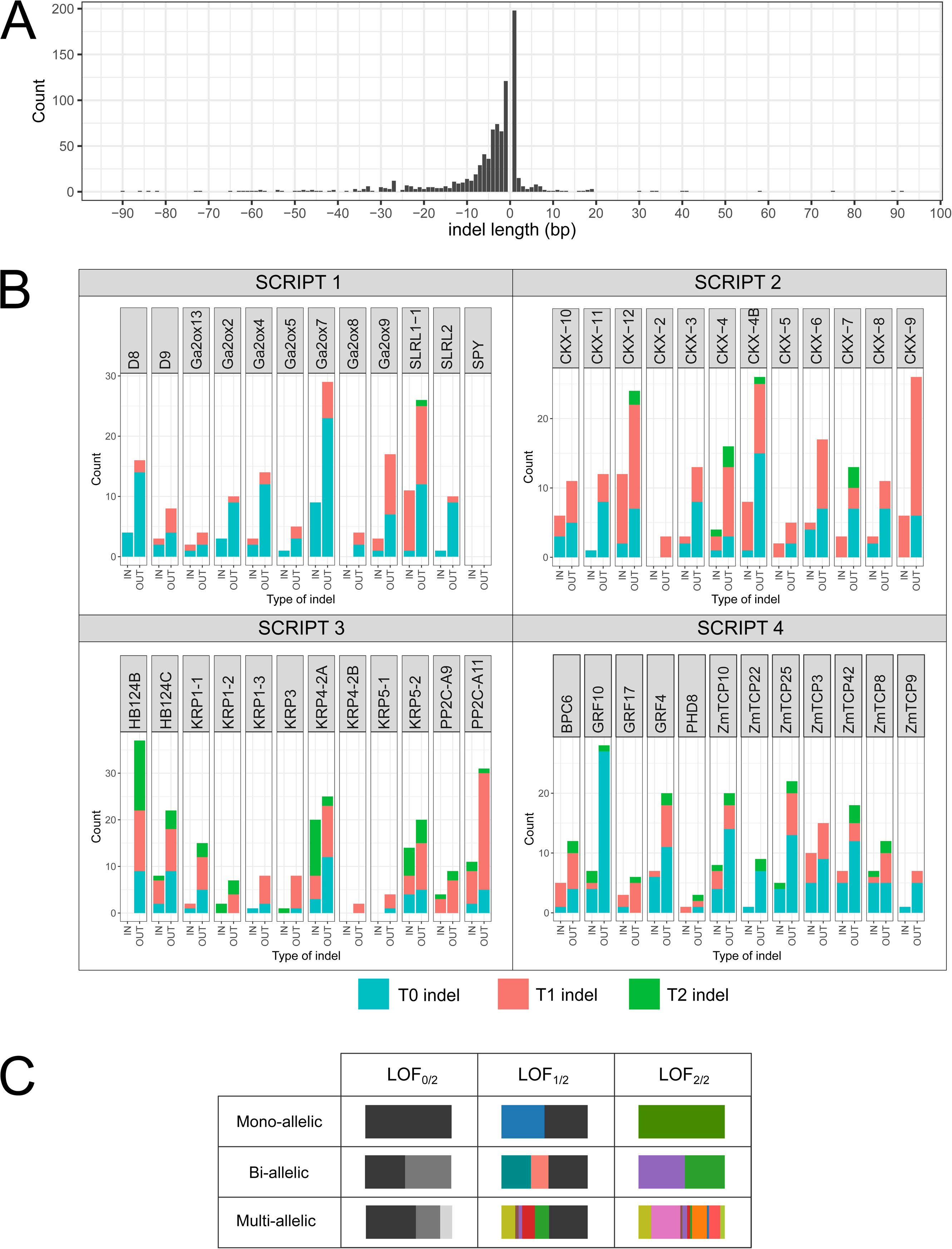
Diversity of mutated haplotypes obtained after CRISPR/Cas9 genome editng. **A**. Distribution of indel length. **B**. Number of different haplotypes with indels observed per gene. Any haplotype with indels with >1% relative frequency in the sequencing reads per locus per sample is included. IN: in-frame indel, OUT: out-frame indel. Blue, red, and green correspond to fractions of indels first observed at T0, T1, and T2, respectively. **C**. Different haplotype combinations in plants can all lead to a gene loss-of-function, either partial (LOF_1/2_) or complete (LOF_2/2_). Each colored horizontal stacked bar corresponds to a different haplotype_KO_. Bar length is proportional to the fraction of sequencing reads per locus containing the haplotype_KO_. The black fraction corresponds to the aggregation of alleles assigned to the wild-type haplotype (haplotype_REF_). For an overview of the different haplotype_KO_ found in T0 plants harboring the different SCRIPTs, see Supplemental Figure S5.

### From haplotype frequencies to genotypic information

The aggregated fraction of haplotype_KO_ in sequencing reads was used as a proxy to characterize partial (LOF_1/2_) and complete (LOF_2/2_) gene KOs (Figure 3). Our approach for the detection of gene edits using HiPlex amplicon sequencing combined with SMAP haplotype-window analyses successfully captured haplotype sequences in 96% of the cases, encouraging us to use this technique to monitor edits in the offspring (Figure 3A). At T0, 73% (35/48) of the target loci showed LOF_1/2_ or LOF_2/2_, with SCRIPT 1 and SCRIPT 3 performing less than SCRIPT 2 and SCRIPT 4 (Figure 3B). At T1, of the 13 remaining genes not edited at T0, 12 (92%) were *de novo* edited. No haplotype_KO_ was observed at the last remaining non-edited locus (*SPY*) at T2. Also, all the transgenerational *de novo* edits were only heterozygous mono-allelic mutations (Figure 3B). Considering both T0 and T1 materials, we observed plants stacking up to nine LOF_1/2_ or LOF_2/2_ in both SCRIPT 1 and SCRIPT 3, and 11 of LOF_1/2_ or LOF_2/2_ gene KOs in SCRIPT 2 and SCRIPT 4 (Figure 3C). Because of sterility issues, progeny of SCRIPT 1 was difficult to generate by crossing, resulting in the low numbers of T2 for that SCRIPT (Figure 3C). Finally, we also studied progeny resulting from inter-script crosses involving two SCRIPTs (2 × 12 target loci) and observed that, on average, 40% of the loci showing edits in the progeny presented transgenerational editing patterns and 25% were completely *de novo* edited, meaning that edits at these loci were not observed in the parental lines (Supplemental Figure S7). Per locus across all populations, on average 7% of the progeny is affected by transgenerational edits inducing LOF_1/2_ at the target sites.

**Figure 3.**
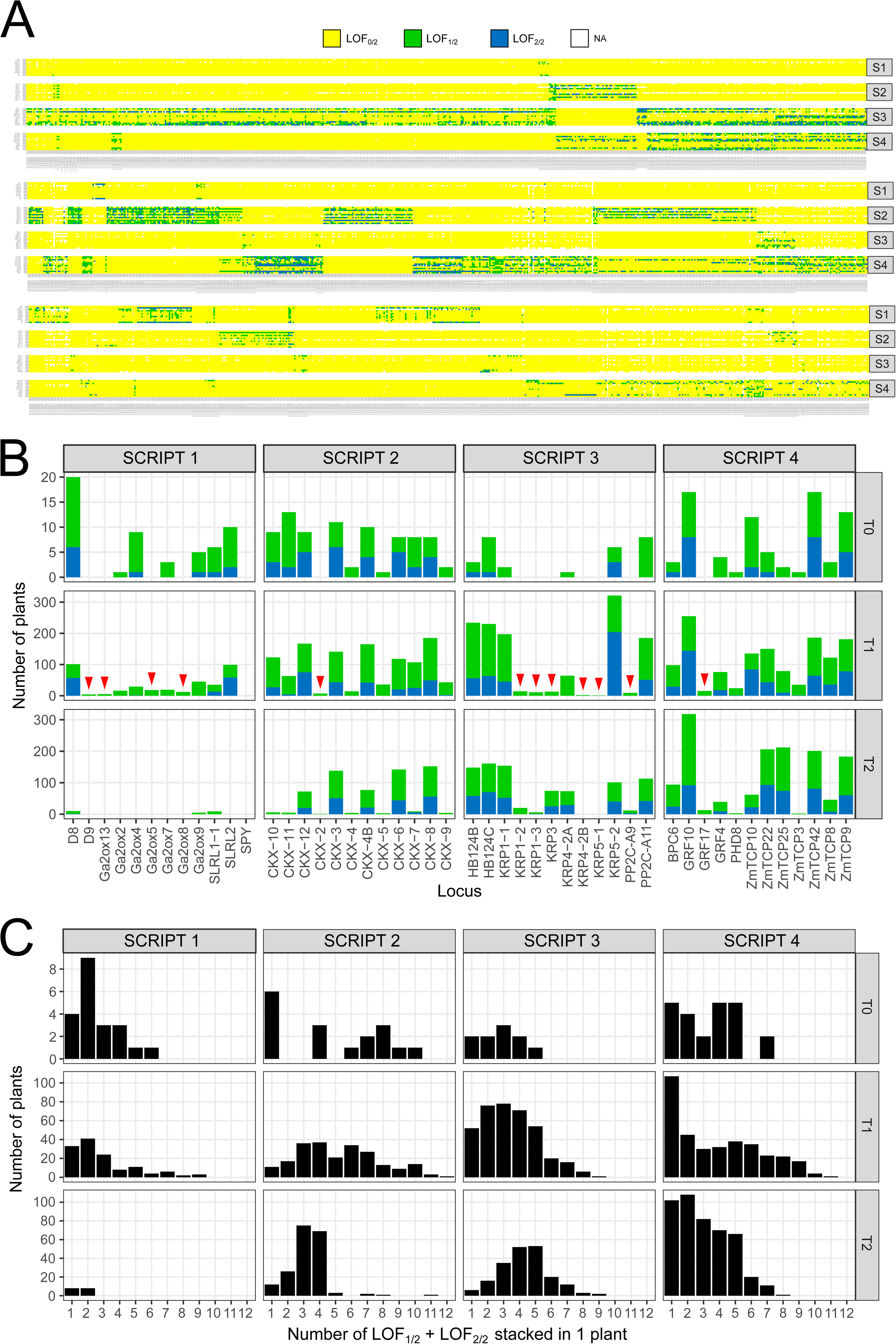
Distribution of LOF in genes across the entire set of samples. Only haplotypeKO were considered for genotype calling. The fractions of reads containing haplotype_KO_ were summed per sample per locus. **A**. Overview of the classes LOF_0/2_, LOF_1/2_, or LOF_2/2_ obtained in the entire sample set for the four SCRIPTS (S1-S4). Samples are on the x-axis and distributed over three rows. **B.** Distributions of LOF_1/2_ (green) and LOF_2/2_ (blue) across the four SCRIPTs throughout generations. The top, middle, and bottom panels show T0, T1, and T2 plants, respectively. Red triangles indicate new LOF that appeared at T1. **C**. Stacking LOF at multiple genes within plants.

In conclusion, the approach of supertransforming EDITOR lines with SCRIPT constructs generated a high frequency of heritable edits in T0 and additional transgenerational edits in T1 and T2.

### T1 single-SCRIPT multiple-edited populations display phenotypic variability in seedling growth-related traits

After we generated the single-SCRIPT populations of edited plants, we studied the effects of multiple gene edits on plant growth by phenotyping T1 maize seedlings derived from T0 selfings of each SCRIPT at the V3 stage. To facilitate high-throughput phenotyping of several populations, we scored easy-to-measure parameters such as the final leaf length and width of leaf 3 (FLL3 and FLW3, respectively) and also integrative parameters such as the fresh weight (FW), dry weight (DW) and moisture content of plants grown under well-watered (WW) and water-deficient (WD) conditions. We scored populations derived from independent transgenic events to analyze the effect of combinations of LOF dosages resulting from different haplotype_KO_ on trait expression (Figure 4, gradient of edits displayed in orange). Detailed information of the different populations that were phenotyped is provided in Supplemental Table S1.

**Figure 4.**
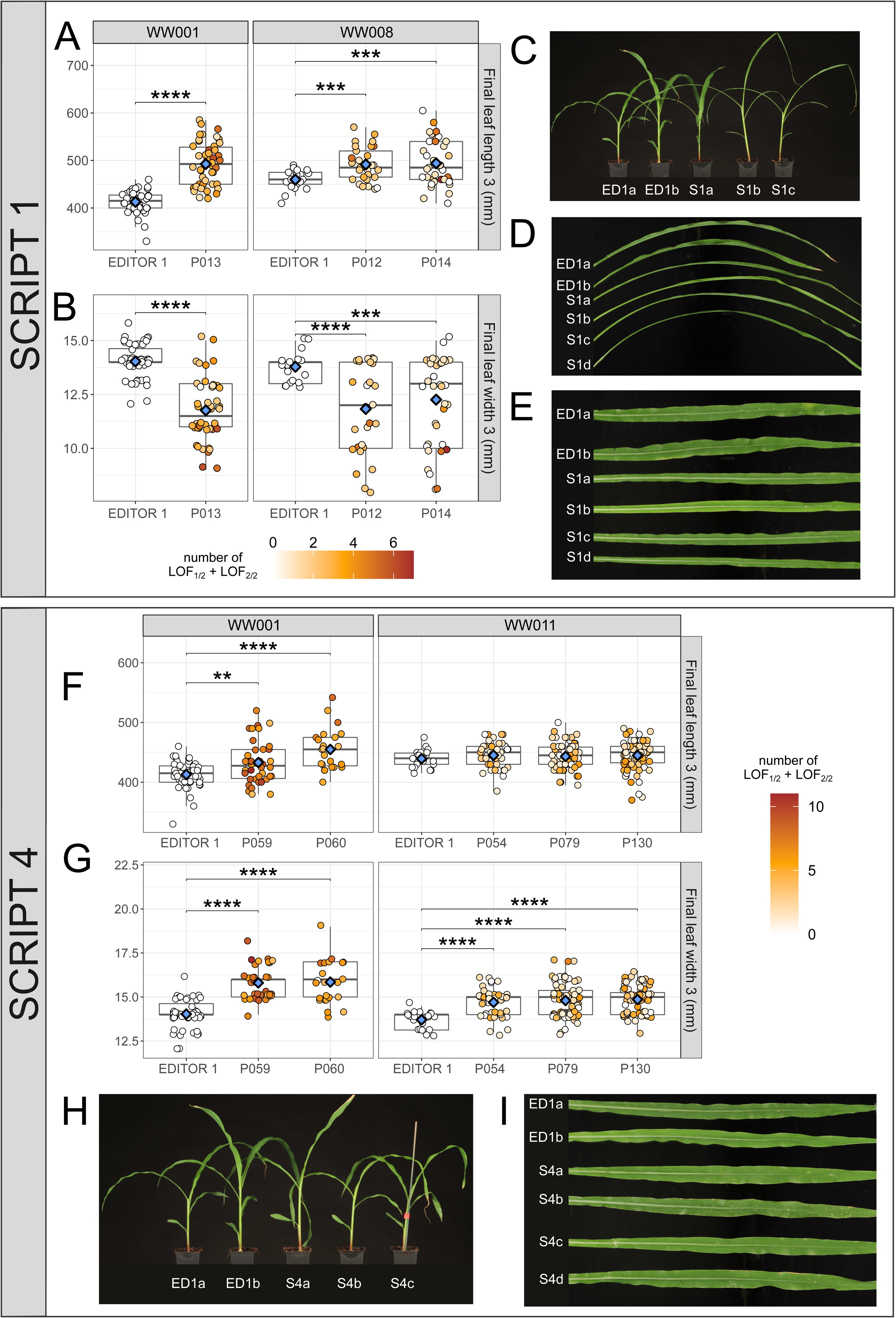
Phenotypes observed in multiple gene-edited populations of SCRIPT 1 and SCRIPT 4. A-B, F-G. Measurements of final length of leaf 3 (FLL3) (A, F) and final leaf width (FLW3) (B, G) of gene-edited SCRIPT 1 (A, B) and SCRIPT 4 (F, G) individuals compared with the EDITOR 1 background control. For each SCRIPT, data corresponds to independent multiple gene-edited populations assayed on two different independent experiments under WW conditions. On the distributions, each dot represents one individual and is colored according to the amount of partial (LOF_1/2_) and complete (LOF_2/2_) LOF observed in that individual. The more orange, the higher the LOF in the individual. Pairwise Student’s t-test were conducted between EDITOR 1 and mutated populations. Significant differences are displayed with p-values summarized as follow: **= p<0.01, *** = p<1e-3, **** = p <1e-4. Blue diamonds indicate the mean of each distribution. C-D, H-I. Pictures of general plant architecture (C for SCRIPT 1 and H for SCRIPT 4) and final leaf 3 (D for SCRIPT 1 and I for SCRIPT 4) compared with the EDITOR 1 (ED1) background.

SCRIPT 1 plants were tested in two independent WW experiments (WW001 and WW008) (Figure 4A, B) and displayed conspicuous phenotypes such as a slender shoot architecture (Figure 4C) with longer and narrower leaves (Figure 4A-B, E) compared with EDITOR 1 controls. The most conspicuous phenotypes could be observed in population P013, which includes individuals with partial or complete LOF in a set of 11/12 genes (Figure 4A-E). Additionally, some SCRIPT 1 plants displayed abnormal tassel development with a lack of florets or pollen and the formation of silks in the anthers (Supplemental Figure S8), leading to male sterility.

For SCRIPT 2 and SCRIPT 3, when tested in experiment WW001, significant increases of about 5% relative to controls could be detected only for FLL3 and only in one of the two populations of each group (P108 for SCRIPT 2, and P033 for SCRIPT 3), while FLW3 remained unaffected in SCRIPT 2 populations or decreased for both populations of SCRIPT 3 (Supplemental Figure S9B-C). Because the genes targeted in SCRIPT 2 are involved in cytokinin metabolism, previously implicated in drought tolerance (Rida et al., 2021), and some of the genes targeted in SCRIPT 3 are drought responsive (Li et al., 2016; Hai et al., 2020), these populations were phenotyped under WD conditions (Supplemental Figure S9B-C). Under WD, the SCRIPT 2 populations showed an enhanced growth (Supplemental Figure S10A), which is reflected by a significant increase in FLL3, FLW3, FW and DW compared with control EDITOR 1 (Supplemental Figure S9B, Supplemental Figure S10 A-C). For SCRIPT 3, all tested populations displayed enhanced growth traits (Supplemental Figure S11), but only significant increases in FLL3 compared with EDITOR 1 were observed (Supplemental Figure S9C). Moreover, population P034 presented a significant increase for FW compared with EDITOR 1 controls (Supplemental Figures S9C and S11).

Changes in leaf morphology were also observed for SCRIPT 4 plants (TCP, GRF family genes). Individuals that segregate LOF dosages in 12/12 and 9/12 genes were observed in populations P059 and P060, respectively (Figure 4F-G, gradient of edits in orange; Supplemental Table S1). Both populations presented significantly longer FLL3 (Figure 4F) alongside a >15% increase in FLW3 compared with EDITOR 1 (Figure 4G-I). The increase in FLL3 could not be detected in populations P054, P079, and P130 (Figure 4F, Supplemental Table S1) but the rise in FLW3 was significantly detected in all populations (Figure 4G).

### Crossing plants with different SCRIPTs allows combining phenotypes in T2 plants

After focusing on single-SCRIPT populations, we phenotyped inter-script populations that stack edits in genes from different SCRIPTs after crossings. For this, T0 plants with different scripts were crossed and the resulting T1 plants (inter-script crosses) were self-crossed. Of all the different combinations, we phenotyped two T2 inter-script populations which presented different profiles of edits in crosses between SCRIPT 2 × SCRIPT 4 (P148 and P152) and SCRIPT 3 × SCRIPT 4 (P157 and P158) under WD conditions. For both populations of SCRIPT 2 × SCRIPT 4 and SCRIPT 3 × SCRIPT 4, we detected a significant increase in FLW3 (Supplemental Figure S12, and Supplemental Table S1), a phenotype observed in single-SCRIPT 4 T1 lines. For the other traits, distinct differences were observed in each population. P148 displayed an increase in FLL3, whereas P152 showed a decrease in FLL3 and significant increases in FW and moisture content compared with the EDITOR 1 control (Supplemental Figure 12A). Both P157 and P158 displayed significant increases in moisture content and P158 displayed reduced DW compared with the EDITOR 1 control (Supplemental Figure 12B).

### Genotype-to-phenotype associations and gene space reduction

After phenotypic evaluation of all SCRIPT populations, we attempted to determine the possible causative genes for the observed phenotypes. Because each individual phenotyping experiment did not allow for sufficient replication of LOF dosages combinations, we performed genotype-to-phenotype associations at the single-gene level. For each gene and trait, we compared the three classes of LOF dosages (LOF_0/2_, LOF_1/2_, and LOF_2/2_) with the EDITOR 1 control. Such single-gene analyses were carried out separately for all experiments conducted under WW and WD, representing in total a collection of more than 1000 plants that include data on selfed, inter- and intra-script crossed lines. Our goal here was to detect major gene effects.

Following this approach, we could detect a subset of genes for each gene family which could be, at least partially, significantly responsible for the observed phenotypes (Figure 5). In SCRIPT 1, phenotypes regarding increases in FLL3 and decreases in FLW3 were associated with edits on DELLA orthologs *D8* and *ZmSLR2* as well as on *ZmGa2ox5* (Figure 5A). For SCRIPT 2, edits in *ZmCKX4B*, *ZmCKX6* and *ZmCKX8* were related to changes in FW, FLW3 and DW (Figure 5B). In the case of SCRIPT 3, LOFs in *ZmKRP5-2* and *ZmPP2C-A11* were associated with increases in FW and DW, while LOF in Zm*KRP1-1*, *ZmHB124B* and *ZmHB124* correlated with increases in biomass moisture (Figure 5C). Finally, for SCRIPT 4, the main genes involved in the increases observed in FLW3 were *ZmTCP8*, *ZmTCP9*, *ZmTCP10*, *ZmTCP22* and *ZmTCP42* (Figure 5D). Particularly, LOFs in *ZmTCP22*, *ZmTCP42* and *ZmTCP9* were associated with concomitant increases in FW and moisture content, and therefore decreases in DW. *ZmGRF10* and *ZmGRF4* were associated to increases in FLL3.

**Figure 5.**
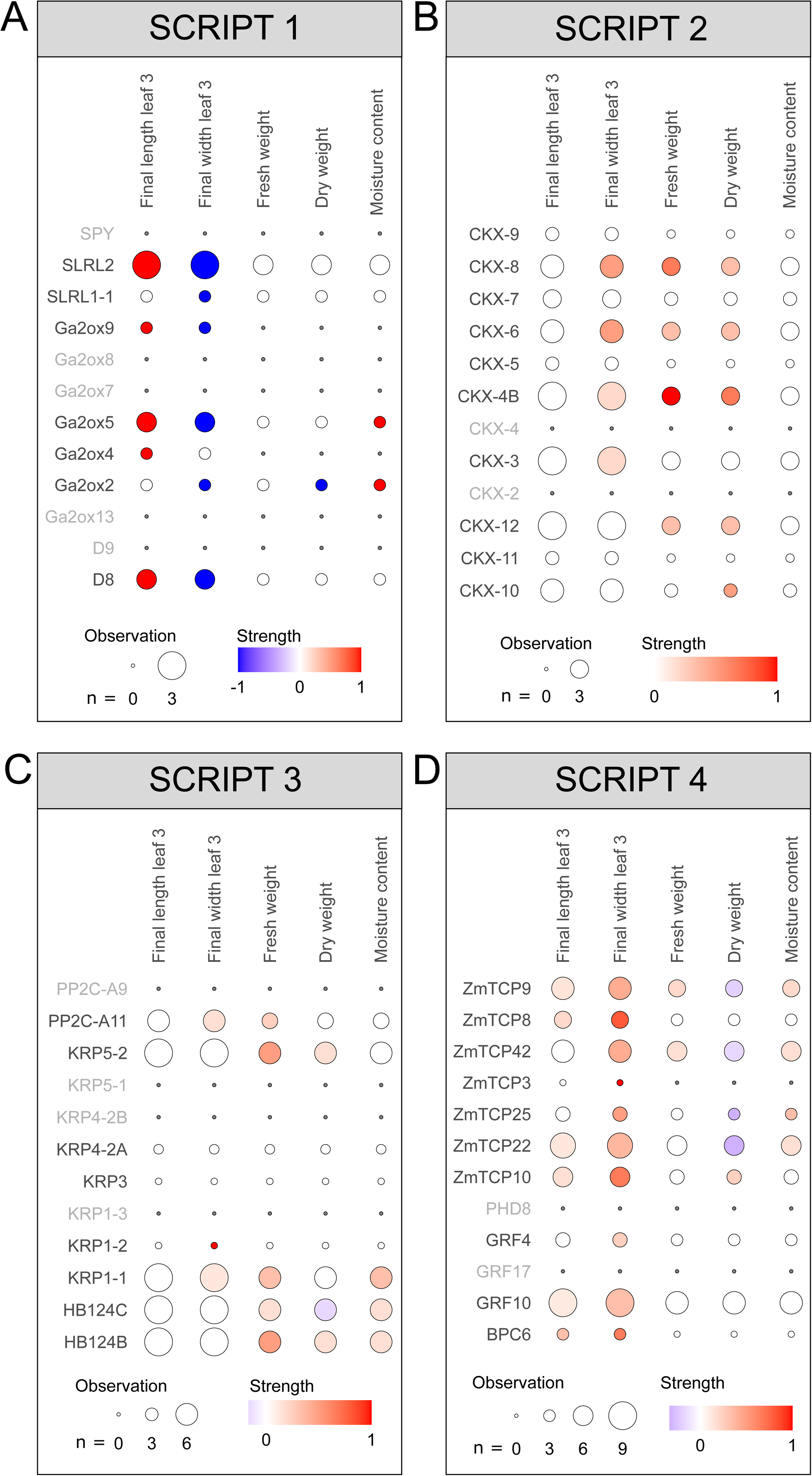
Aggregated association analysis of single-gene LOF and traits. Summaries of single-gene associations to traits are represented for SCRIPT 1 (A), SCRIPT 2 (B), SCRIPT 3 (C), and SCRIPT 4 (D). Single-gene associations were performed per population, in each phenotypic experiment and for all measured traits. Results are summarized per gene, per trait with two indices. 1) Observation: the number of time a given gene has been observed in a situation with sufficient genotypic and phenotypic data across populations and experiments. An observation with sufficient data corresponds to a situation where a gene displays at least one LOF group between LOF_1/2_ and LOF_2/2_ represented by at least six individuals with phenotypic information for a specific trait. In such cases, the mean phenotypic value of each genotypic group could be statistically compared to the mean phenotypic value of the EDITOR 1 control. 2) Strength: for each gene, we calculated the weighted sum of observations in which the genotypic group with the highest mean phenotypic value is above (weight: +1) or below (weight: -1) 10% the mean phenotypic value of EDITOR 1. The resulting sum was divided by the total number of observations (n). Associations displaying highest strength, either positive or negative, along with a large total number of observations indicate strong evidence for a gene effect on the trait.

To further validate the rationale used for the associations, we analyzed in detail population P012 of SCRIPT 1 (Figure 6). In this population, *D8*, one of the selected genes associated to increases in FLL3 (Figure 5A), showed two haplotype_KO_, each with an out-frame indel (-1 bp and +1 bp) and an haplotype_REF_ with an in-frame indel (-3 bp) (Figure 6A). In the progeny, the haplotypes segregated resulting in different LOF dosage combinations. Within that population, plants containing only a LOF_1/2_ in *D8* presented similar phenotypes of FLL3 and FLW3 compared with EDITOR 1, whereas plants with a LOF_2/2_ in *D8* displayed longer and narrower leaves (Figure 6B-C).

**Figure 6.**
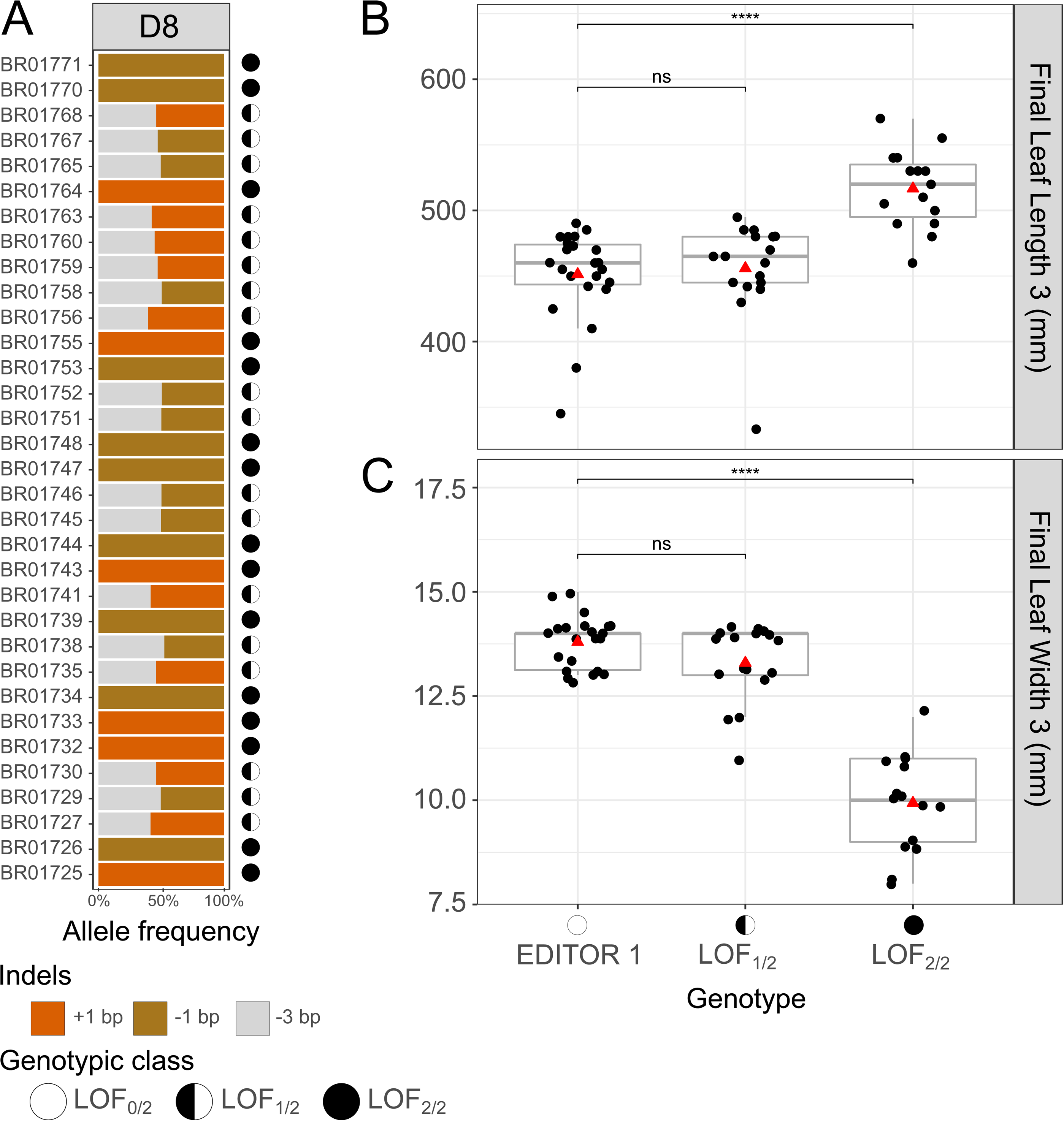
LOF dosages in *D8* and leaf shape parameters. **A**. Haplotype profiles at gene *D8* of T1 segregants from population P012. Three haplotypes were detected, with two containing out-frame indels (-1 bp, brown and +1 bp, orange) and one containing an in-frame (-3 bp, gray) deletion. This results in a collection of plants with *D8* either partially (LOF _1/2_) or completely (LOF_2/2_) knocked out. The resulting two classes of LOF dosages are compared to EDITOR 1 for final leaf length 3 (**B**) and final leaf width 3 (**C**). Significant differences (pairwise Student’s *t*-test) are displayed with p-values summarized as follow: *** = p<1e-3, ns: not-significant. Red triangles indicate the mean of each distribution.

## Discussion

Complex agronomical traits such as yield or resistance to a particular (a)biotic stress are governed by a large network of genes that together determine a specific phenotype. Understanding the complexity of such networks is the central aspiration of systems biology.

Here, we developed an experimental approach, named BREEDIT, to study gene networks affecting complex quantitative traits by combining multiplex CRISPR-mediated gene editing of whole gene families with specific crossing schemes. In BREEDIT, a Cas9-expressing line (EDITOR) is supertransformed with vectors containing 12 gRNAs (SCRIPTs) targeting a set of GRGs. Gene edits are further stacked in plants using crossing schemes.

We evaluated the BREEDIT strategy by targeting putative players in major plant gene families or pathways involved in growth regulation. The success rate of the multiplex gene editing approach in maize was very high, with more than 97% of the genes showing at least partial or complete LOF at T1. In just two generations, BREEDIT created multiple gene KOs leading to a diverse collection of genetic profiles, from low-order mutants with one, two or three gene KOs to higher-order mutants stacking mutations on up to 11 genes out of the 12 within a single SCRIPT. Additional levers could be used to further increase the number of gene KOs stacked in one plant, namely inter-script crosses and the ongoing Cas9 activity. We indeed showed that higher-order mutants could be obtained by crossing plants already containing high numbers of gene KOs at the single-SCRIPT level. Finally, the combined presence of the Cas9 and the SCRIPT throughout the generations enabled the CRISPR/Cas9 machinery to continuously generate mutations in not yet edited loci. This ongoing Cas9 activity can therefore contribute to reach saturation in gene edits after a couple of generations and increase the number of gene KOs stacked in one plant. Interestingly, we noticed that while in the T0 plants often both copies of the target genes carried the same or a different mutation (bi-allelic), genes newly edited at T1 all showed heterozygous mutations, suggesting that only one chromosome of the two is edited. We hypothesized that chromatin condensation may influence DNA accessibility for the CRISPR/Cas9 machinery, possibly by imprinting (Borg and Berger, 2015). Further research is needed to elaborate on the mechanisms.

We obtained more than 1000 plants with often different unique LOF profiles to score for phenotypes. The high sensivity of HiPlex amplicon sequencing enabled to capture complete sets of haplotypes with CRISPR/Cas9 mutations in large arrays of samples and loci. We used the haplotype sequence to focus on haplotypes supposed to have major effect (haplotype_KO_) on the translated protein function or activity. The experimental set-up based on multiple observations of significant single-gene KO associations with phenotypes across different populations and experiments enabled us to identify significant phenotypic responses in growth traits for all SCRIPTs. In the case of SCRIPT 1, we observed previously known effects of elevated gibberellic acid (GA) levels, such as plants with long and narrow leaves (Nelissen et al., 2012; Voorend et al., 2016) as well as male sterility, a trait that was previously associated with the effect of GA on tassel development (Colombo and Favret, 1996). SCRIPT 4 plants displayed an increased FLW3 (alongside with milder increases in FLL3), which impacted the FW in some populations. Lastly, for SCRIPT 2 and SCRIPT 3, the most pronounced phenotypes observed were increased FLL3 and FW, particularly under WD conditions.

If we consider these single-SCRIPT lines as building blocks, the possibility to stack several different combinations of SCRIPTs by crossing, opens the way to create higher-order mutant lines that may display even stronger or additive phenotypes. The inter-script lines we generated displayed higher-order mutations more than (12) and inherited traits observed in parental single-SCRIPT lines. Though some of the expressed traits (e.g. increases in FW) were not observed in all different inter-script populations of the same type (probably because different haplotypes are segregating in different populations), some other traits (such as increases of FLW) were consistently observed in all generations of lines containing SCRIPT 4, which further validates the consistency of the results analyzed at single-SCRIPT level.

Another important outcome of the BREEDIT strategy is the possibility to screen large sets of genes that are then ranked and prioritized to delineate a minimal set of LOF required to induce a maximal phenotypic effect. Further inspection of the selected 48 GRGs showed that certain subsets of genes are strongly associated to specific traits (or combination of traits). Using this information, we built a possible regulatory network that integrates all the single-gene effects and their impact over all the different measured traits (Figure 7). In this network, central genes (such as *ZmCKX4B, ZmCKX48 and ZmCKX46* and *ZmTCP9*, *ZmTCP10*, *ZmTCP22* and *ZmTCP42*) act as nodes connected to several traits, inferring a possible broader role in regulation. The genes located at the edge of the network may play a more defined role, connecting to just one or two traits (such as *ZmGRF10*, *ZmGRF4, ZmCKX3* and *ZmTCP8*). Finally, other genes connect to specific patterns, like *D8*, *SLRL2*, and *Ga2ox5*, of which LOF strongly increases the FLL3 while also strongly decreasing the FLW, or *ZmHB124B*, of which LOF influences the FW and moisture positively. Therefore, using single-gene KO associations, we could identify subsets of genes per family, of which LOF, alone or in combination, may be responsible for the observed phenotypes. Furthermore, some of these genes were already shown to play roles in modifying agronomical traits in other studies. A gain-of-function mutation in *D8* (SCRIPT 1), encoding a DELLA maize protein ortholog, is causative of dwarf phenotypes (Winkler and Freeling, 1994; Lawit et al., 2010). Zm*CKX-4B* (SCRIPT 2) plays a role in both drought (Rida et al., 2021) and heat shock stress (Wang et al., 2020). Downregulation of *ZmHB124B* and *ZmHB124C* induces the formation of additional protoxylem files in the vasculature (Bloch et al., 2019), which could prevent vascular embolism and water retention under water-limiting conditions (Hwang et al., 2016). *ZmGRF10* (SCRIPT 4) overexpression in maize leads to a decrease in leaf length and height (Wu et al., 2014; Nelissen et al., 2015). Although these genes were highlighted in our single-gene KO associations, we cannot exclude that other genes from the original pool may play minor roles, either individually or in combinations, but end up masked by the effect of major gene KOs. Nonetheless, this subset selection provides a valuable material pool for further research to tackle specific traits (e.g. leaf width, enhanced growth under WD).

**Figure 7.**
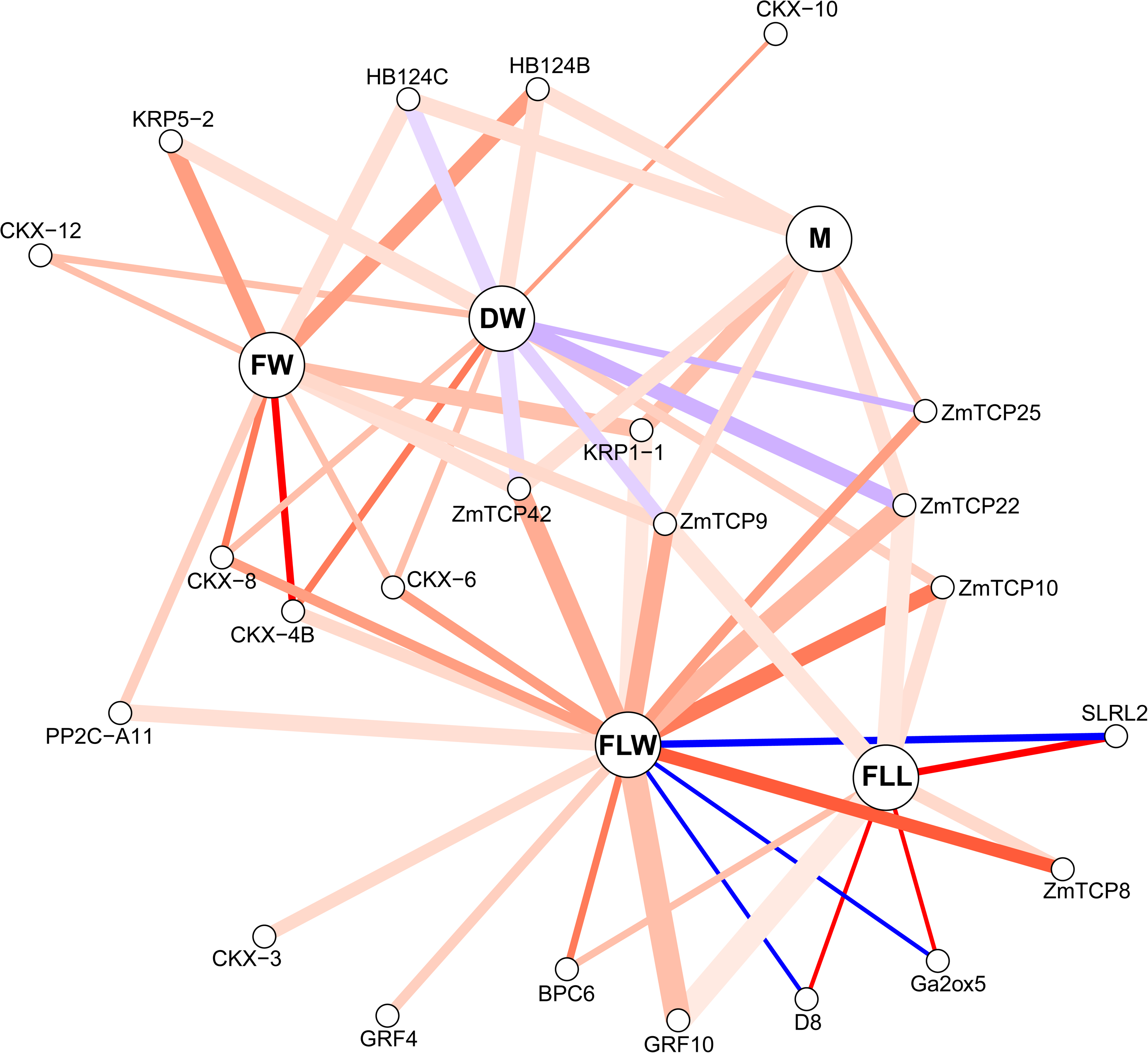
Network representation of single-gene effects on growth-related traits. Traits are displayed in bold (FLL: final leaf length; FLW: final leaf width; FW: fresh weight; DW: dry weight; M: moisture content). Genes at least associated once with a trait are displayed. Lines indicate connections between genes and traits. Line width is proportional to the number of times the underlying dataset to detect a gene knockout-trait association in different experiments and/or populations contained sufficient data for statistics (i.e., minimum one LOF class between LOF_1/2_ and LOF_2/2_ with at least six individuals with phenotypic information). Line color represents the weighted fraction of gene KO-trait associations that significantly outperformed by 10% the EDITOR 1 control (ANOVA test; p<5%), either positively (weight: +1, more red) or negatively (weight: -1, more blue), over the number of times a gene KO-trait association could have been observed because of sufficient data points.

While applying the BREEDIT strategy to our case study in maize, we identified a couple of limits to the approach. First, the inability to fully uncouple complex gene interactions. In plant models where transformation and regeneration are efficient, the possibility for massive gene editing grows faster than the capacity to phenotypically analyze the resulting collections of mutants. For complex quantitative traits, large populations of lower-order mutants need to be screened accurately to decipher the complex mechanisms underlying plant development (Liu et al., 2020). To illustrate this, we have developed the following multiplex edited scenario (Supplemental Figure S13). If one would be interested in exhaustively capturing both additive and synergistic gene effects, all the gene KO combinations have to be generated and analyzed. Considering n genes to be targeted, the number of different genetic combinations that have to be produced amounts to 2^n^ in the case of two-state genes (homozygous wild type or homozygous mutant) (Supplemental Figure S13A) and 3^n^ if the heterozygous stage has to be considered as well. Given that at least ten replicates per genetic profile (combination) are required to statistically demonstrate a 10% significant difference in FLL3 with enough statistical power (Supplemental Figure S13B), the final amount of plants to be processed increases dramatically as the number of genes in the study grows. Large scale phenotyping/genotyping in field conditions would allow to increase the statistical power to detect combinatorial genes effects that govern agronomic traits, including seed yield.

By applying the BREEDIT strategy on a broad gene set, one could generate a reduced sub-selection of genes of interest underlying a trait and then apply complementary approaches to rule out their contribution to a specific trait of interest in just two generations. One of such approaches is the use of haploid induction (Chaikam et al., 2019; Jacquier et al., 2020), a promising technique to create homozygous mutations, thus removing the need to consider heterozygous material. This could be particularly interesting as a follow up of BREEDIT, because the new gene edits that we observed at T1 were all heterozygous. Another approach is to preselect plants to be phenotyped on the basis of their genetic constitution by predicting gene effects with statistical models in the same fashion as for genomic selection. Such predictions can be combined with the use of non-destructive seed chipping (Mills et al., 2020) to pick specific gene combinations before sowing and therefore reduce the number of plants to be tested. Once the gene space is lowered, the BREEDIT pipeline can be followed again to design validation constructs by engineering a vector containing gRNAs targeting only the genes retained in the selected subset.

In this study, we present the BREEDIT strategy to rapidly generate a large collection of mutants in specific gene families, pathways or networks. We foresee a large potential for BREEDIT combined with existing and more recent breeding approaches, such as marker-assisted breeding, haploid induction, and genomic selection. Effectively implementing the concept of breeding by editing using the BREEDIT pipeline still requires to overcome some practical obstacles, such as the ability to transform and regenerate the plant material, obtain the desired gene KOs and segregate out the original transgene construct. When these conditions are met, applying the BREEDIT pipeline enables to generate many lines with specific combinations of gene KOs able to modify particular traits of interest. These engineered lines could be directly introduced in a hybrid breeding pipeline by crosses to elite material. The impact of favorable allele combinations on complex traits could furthermore be evaluated in different genetic backgrounds and across several generations to assess their heritability. In that perspective, BREEDIT could significantly speed up pre-breeding activities in which pools of diverse materials (wild species, landraces, commercial varieties) are usually screened for promising mutations and phenotypes (Teixeira and Guimarães, 2021) that then have to be transferred into an intermediate set of materials that breeders can use to create new varieties. Introgression of alleles from a divergent pool of materials is often cumbersome because of cross incompatibility, low seed yield quantity and quality, or persistence of a deleterious linkage drag. Provided that elite materials can be transformed and regenerated, the reverse-genetics approach developed in the BREEDIT pipeline can circumvent the long and tedious step of introgression and save time in the development of new commercial varieties. An additional benefit of BREEDIT could be the generation of large collections of plants mutated in coding or non-coding genome areas using other novel CRISPR technologies such as base editing and promoter bashing to further extend the repertoire of allele variability and phenotypic responses (Vats et al., 2019; Anzalone et al., 2020; Gaillochet et al., 2021).

## Materials and methods

### Plant material and DNA extraction

The original line used for all transformation procedures was the maize inbred line B104. DNA was extracted following an adapted protocol from Berendzen et al. (2005) coupled with a magnetic beads purification. A piece of 1-2 cm of leaf 1 was ground in 8-strip 2-ml capacity tubes (National Scientific Supply Co). After grinding and centrifugation, the supernatant was mixed with magnetic beads (CleanNA), washed in 80% ethanol and dried for further processing.

### Selection of GRGs and curation of gene models

We selected 48 GRGs based on the literature, in house knowledge and orthology searches (see results) in version 4 of the reference maize B73 genome sequence (Jiao et al., 2017). The integrative orthology viewer in PLAZA v4.5 (https://bioinformatics.psb.ugent.be/plaza/versions/plaza_v4_5_monocots/) was used for identification of most orthologous genes, both for finding gene families from other species or for identification of the corresponding maize B104 gene IDs. When required, B104 gene models were manually curated using ORCAE, an online genome annotation resource (https://bioinformatics.psb.ugent.be/orcae/). Comparison of sequences in maize lines B104 and B73 was done by pairwise alignment using Geneious Prime 2020.1.2 (https://www.geneious.com/prime/). Design of the amplicons and gRNAs was performed in Geneious Prime. The maize B73 genome version 4 was used to identify gRNA on-target and off-target sites. gRNAs were selected with specificity score ≥80-85%, no stretch of Ts (>4), without internal *Bsa*I or *Bbs*I restriction sites, which would interfere gRNA expression and/or vector construction, respectively.

### Monitoring CRISPR/Cas9 edits with HiPlex amplicon sequencing

Geneious Prime was used to design primers to amplify the genomic regions surrounding the gRNA cutting sites. A manual selection of two amplicons per gene was done with a size range between 120-150 bp. Each amplicon contained at least two gRNAs separated from either primer by at least 15 bp. The amplicons were selected to target the middle of the coding sequence and to not overlap. The specificity of primers was checked in the maize B73 genome version 4 and only specific primers were retained (Supplemental Table S2). All primers were pooled in a HiPlex amplicon sequencing assay to sequence each locus in each plant simultaneously. HiPlex library preparation was performed by Floodlight Genomics facility (Knoxville, TN, USA) using the MonsterPlex technology. Pilot runs of HiPlex amplicon sequencing were conducted to select the best amplicon per gene (out of the two). The selection was based on amplification efficiency in the HiPlex assay measured as read counts and unambiguous read-reference mapping. Per gene, the overlapping gRNA was selected for cloning into the expression vector. We used SMAP haplotype-window (Schaumont et al., 2022) to trim sequencing reads, identify haplotypes at each locus, and calculate the respective haplotype frequency per locus per sample. SMAP haplotype-window extracts haplotypes from the HiPlex sequencing reads as the entire DNA sequence between the HiPlex primers per locus. Any unique DNA sequence is considered as a haplotype. The total haplotype count is recorded per locus per sample and the relative haplotype frequency per locus per sample is calculated. A haplotype detection threshold of at least 1% relative read depth per locus per sample was set to remove possible spurious haplotypes derived from amplification and/or sequencing artifacts. The nucleotide length difference between the haplotype sequence and the B104 reference sequence (LDR) was used to classify the mutations into three classes: SNPs (LDR = 0 but sequences are different), insertions (LDR > 0, the mutated haplotype is longer than the reference haplotype), and deletions (LDR < 0, the mutated haplotype is shorter than the reference haplotype). We defined haplotypes whose indel length is not a multiple of three nucleotides as haplotype_KO_ because they generate a frame shift in the open reading frame that likely leads to the translation of a wrong amino acid sequence downstream of the mutation, and/or create a premature stop codon, both of which could disrupt the protein function or activity.

Haplotypes with SNPs outside the cutting site and in-frame indels are referred as to Haplotype_REF_ to denote possible minor impact of their mutations on the resulting protein, which may still behave as the reference protein.

Maize is a diploid organism in which each gene has two alleles per nucleus, each derived from one of the two parents. In plant material that stably express CRISPR/Cas9 and gRNAs, continuously driving gene editing, one may expect to observe mosaic tissues, i.e. patches of tissues within an organ that contain different genome sequences due to non-uniform gene editing. Mosaic tissues may occur both in primary transformants and subsequent generations because of the initial and ongoing Cas9 activity, respectively. A single leaf sample used for DNA preparation may therefore contain cells with different gene edits resulting in scoring one individual with more than two alleles. The allele dosage is also affected by mosaicism. Multi-allelism resulting from mosaic tissues blurs the expected 50:50 read depth ratio commonly observed between the two alleles of a diploid organism. In addition, bi-allelism can be observed in non-mosaic tissues, with a plant harboring two indels of a same or different nature (in-frame or out-frame) following a 50:50 read depth ratio. Genotype-to-phenotype statistical associations require discrete genotypic classes (absence/presence, or homozygous wild-type, heterozygous, homozygous mutant). We therefore summed the relative fraction of haplotype_KO_ per locus per sample to quantify how much the locus is affected by mutations leading to a LOF. The resulting aggregation (Σhaplotype_KO_) is discretized in three genotypic classes representing three dosages of haplotype_KO_: LOF_0/2_ (<15% of the read depth per locus per sample contain haplotype_KO_), LOF_1/2_ (between 40% and 60% of the read depth per locus per sample contain haplotype_KO_), and LOF_2/2_ (>85% of the read depth per locus per sample contain haplotype_KO_). Because distinguishing between these three groups is critical to analyze dosage effects associated with a particular trait, any value outside of these three ranges was scored as missing genotype call during the genotype-to-phenotype association analyses.

### Construction and cloning of SCRIPT vectors

The gRNA entry vectors were constructed by PCR amplification with Q5 High-Fidelity DNA polymerase (New England Biolabs) of the entire pGG-[B-F]-OsU3-BbsI-ccdB-BbsI-[C-G] plasmids according to the manufacturer’s guidelines. The primers contained an extension to insert unique linkers (Torella et al., 2014) between the scaffold and OsU3 promoter (Supplemental Table S3 and Supplemental Table S4). Two of the five linkers were modified to contain *Not*I restriction sites to facilitate validation of the final expression vectors by restriction digest (Supplemental Figure S1). Gibson assembly was performed with NEBuilder Hifi DNA Assembly Mix (New England Biolabs) to circularize the PCR products into entry vectors using the manufacturer’s guidelines. The new entry vectors were confirmed by Sanger sequencing (Mix2Seq service, Eurofins Scientific).

The gRNA construction and Golden Gate assembly into binary vectors was done as previously described (Decaestecker et al., 2019). Briefly, paired gRNA entry vectors were created by PCR amplification (Red Taq DNA Polymerase Master Mix, VWR Life Science or iProof High-Fidelity DNA Polymerase, Bio-Rad Laboratories) on the template plasmid pEN-2xTaU3 with primers containing the 20-nt spacer sequences and *Bbs*I restriction sites. Column-purified PCR products were cloned into the Golden Gate entry vectors via a Golden Gate reaction using *Bbs*I (New England Biolabs). All paired gRNA entry vectors were verified by Sanger sequencing.

Expression vectors (SCRIPT 1-4; Supplemental Figure S1) were constructed by a Golden Gate reaction with *Bsa*I (New England Biolabs) using the paired gRNA entry vectors and a destination vector as previously described (Decaestecker et al., 2019). The destination vector, pGGBb-AG, contains a GreenGate destination module (AG) and a bialaphos-resistance (*bar*) gene driven by the 35S promoter. The expression of each individual gRNA was alternatively driven by either the rice OsU3 promoter or the wheat TaU3 promoter (Xing et al., 2014). The SCRIPT vectors were transformed via heat-shock into *ccd*B-sensitive DH5α *Escherichia coli* cells, grown on LB medium containing 100 µg/mL spectinomycin and extracted with the GeneJET Plasmid Miniprep Kit (Thermo Fisher Scientific). Quality control was performed by restriction digest with *Not*I (Promega). SCRIPTs were transformed into *Agrobacterium tumefaciens* EHA 105 cells by freeze/thaw method and plated on YEB medium with 100 µg/mL rifampicin and 100 µg/mL spectinomycin. The gRNA entry and pGGBb-AG destination vectors can be obtained via https://gatewayvectors.vib.be/.

### Generation of EDITOR maize lines

The *zCas9* coding sequence containing a *Zea may*s-codon optimized Cas9 (Xing et al., 2014) was cloned under control of the *ZmUbi1* promoter (pZmUBIL) and NOS terminator in pEN-L4-AG-R1 (Houbaert et al., 2018) using GreenGate cloning (Lampropoulos et al., 2013). The transcriptional unit was recombined with pEN-L1-linker-L2 and the pHbm42GW7 destination vector (Karimi et al., 2013). The resulting construct pXHb-pZmUBIL-zCas9-NOSt allows to select maize transformants with hygromycin and is referred to as the EDITOR construct.

The EDITOR construct was introduced into the B104 maize line using *Agrobacterium*-mediated transformation of immature embryos (Coussens et al., 2012) and hygromycin as a selection agent. Three independent lines (EDITOR 1 to 3) with a single-locus insertion event were selected and made homozygous for the T-DNA locus by self-crossing. To measure Cas9 protein levels, total proteins from leaf tissue of EDITOR lines were extracted, separated by polyacrylamide gel electrophoresis and blotted on PVDF membrane. For quantification, the blots were stained by incubation with anti-Cas9-HRP primary antibody (Abcam, 1:5000) for 4 h and detected by chemiluminescence. Blots were also Ponceau-stained for protein loading control. EDITOR 1 was crossed with wild-type B104 plants to yield heterozygous immature embryos for a second round of transformation (supertransformation) with each SCRIPT construct separately. Backcrosses render more seeds/embryos compared to self-crosses and also facilitate the removal of Cas9 in the progeny by segregation. For each SCRIPT, at least ten independent T0 supertransformants following BASTA selection were obtained and genotyped by HiPlex amplicon sequencing.

### Experimental design and phenotyping

Maize seeds were soaked in water for 24 h and then sown in 0.3-l square pots (7×7×7 cm) using ‘potgrond met meststof’ (N.V. Van Israel) as substrate. Pots were then arrayed in groups of 24 in 48.0-x 30.5-cm trays, randomized and placed in growth chambers with controlled temperature (24°C), relative humidity (55%) and a 16:8 photoperiod with controlled light intensity (170-200 µmol/m²/s photosynthetic active radiation provided from a mixture of 50/50 Radium halogen HRI-BT 400W/D Pro Daylight and Philips master son-t pia plus 400-W bulbs).

For WW conditions, plants were grown under a water regime of 2.4 g of water per g of dry potting mix, while for WD assays, this was reduced to 1.1 - 1.4 g of water per g of dry potting mix, which is approximately -100 kPa water potential (Verbraeken et al., 2021). The final leaf length was measured at V3 (FLL3, when the collar of leaf 3 is fully developed) from the crown of the plant to the leaf tip and final leaf width (FLW3) was determined at the middle point of the leaf blade. For biomass, aerial parts of V3 seedlings were harvested and weighed for fresh weight (FW) and then dried in a 60°C oven to estimate dry weight (DW). Biomass moisture content was calculated on a DW basis as FW-DW/DW.

### Statistics to detect genotype-to-phenotype associations

Phenotypic datasets were trimmed to remove individuals that scored as under-developed (misshapen or developed less than V3 at the moment of harvest, or over-grown, i.e. that surpassed V3 at the moment of sampling) during the phenotyping trials. Within each population and experiment, one-way ANOVAs were then conducted at the single-gene level to check for differences between the control (EDITOR 1) and mutant groups (either LOF_1/2_ or LOF_2/2_). The minimal size of a mutant group to be considered in statistical analysis was six individuals having both phenotypic and genotypic data. Post-hoc HSD Tukey’s tests were then performed to assign each mutant group to a statistical group. Finally, we recorded the number of times a KO (either LOF_1/2_ or LOF_2/2_) of a specific gene was found to be significantly associated with a given trait while leading on average to a >10% increase or decrease compared to the control line (EDITOR 1). We compared that count with the number of times sufficient data was available to statistically conclude on a gene KO effect and defined the resulting ratio as the strength of the association.

### Accession numbers

The entire set of Illumina paired-end read sequences have been deposited at the Sequence Read Archive (DDBJ/ENA/GenBank) under the BioProject accession PRJNA815957.

## Supplemental data

**Supplemental Figure S1.** Map of a SCRIPT construct containing 12 gRNAs.

**Supplemental Figure S2.** Location of the 48 growth-related genes over the 10 chromosomes of maize.

**Supplemental Figure S3.** Cas9 protein expression levels in three different EDITOR lines.

**Supplemental Figure S4.** Comparative summaries of the mutations observed in plants harboring SCRIPT 1 in an EDITOR 1 vs. EDITOR 3 background.

**Supplemental Figure S5.** Haplotype overview in T0 material from the four SCRIPTs.

**Supplemental Figure S6.** Distribution of the aggregated frequency of haplotypes with out-frame indels (Σ Haplotype KO) in the ingroup samples.

**Supplemental Figure S7.** Appearance of new indels in inter-script crosses.

**Supplemental Figure S8.** Phenotypes of mutated populations for the four SCRIPTs.

**Supplemental Figure S9.** Sterility phenotypes observed in tassels of SCRIPT 1 T0 plants.

**Supplemental Figure S10.** Characteristic phenotypes observed in T1 plants of SCRIPT 2 under water-deficient conditions.

**Supplemental Figure S11.** Characteristic phenotypes observed in T1 plants of SCRIPT 3 under water-deficient conditions.

**Supplemental Figure S12.** Phenotypes of mutated inter-script populations.

**Supplemental Figure S13.** Considerations on numbers for multiplex gene editing experiments.

**Supplemental Table 1:** Detailed information of populations used in the phenotyping assays. The number of loci edited correspond to segregating or fixed loci with an aggregated ratio of indels of 40% in at least 3 plants in the population.

| ID population | Scripts | Generation | Type | Total edited loci | Fixed loci | Experiment |
| --- | --- | --- | --- | --- | --- | --- |
| P012 | S1 | T1 | Self | 9/12 | 1 | WW08 |
| P013 | S1 | T1 | Intrascript | 11/12 | 2 | WW01 |
| P014 | S1 | T1 | Self | 9/12 | 1 | WW08 |
| P019 | S2 | T1 | Self | 10/12 | 2 | WW01, WD03 |
| P108 | S2 | T1 | Intrascript | 11/12 | 0 | WW01, WD05 |
| P032 | S3 | T1 | Self | 6/12 | 0 | WW1 |
| P033 | S3 | T1 | Self | 7/12 | 1 | WW1,WD07 |
| P034 | S3 | T1 | Intrascript | 10/12 | 1 | WD07 |
| P030 | S3 | T1 | Self | 5/12 | 0 | WD07 |
| P059 | S4 | T1 | Intrascript | 12/12 | 5 | WW01 |
| P060 | S4 | T1 | Self | 9/12 | 5 | WW01 |
| P054 | S4 | T1 | Self | 9/12 | 0 | WW11 |
| P079 | S4 | T1 | Self | 10/12 | 0 | WW11 |
| P130 | S4 | T2 | Self | 11/12 | 0 | WW11 |
| P148 | S2 + S4 | T2 | Interscript | 7/24 | 0 | WD09 |
| P152 | S2 + S4 | T2 | Interscript | 8/24 | 0 | WD09 |
| P157 | S3 + S4 | T2 | Interscript | 16/24 | 0 | WD06 |
| P158 | S3 + S4 | T2 | Interscript | 16/24 | 0 | WD06 |

**Supplemental Table 2:**
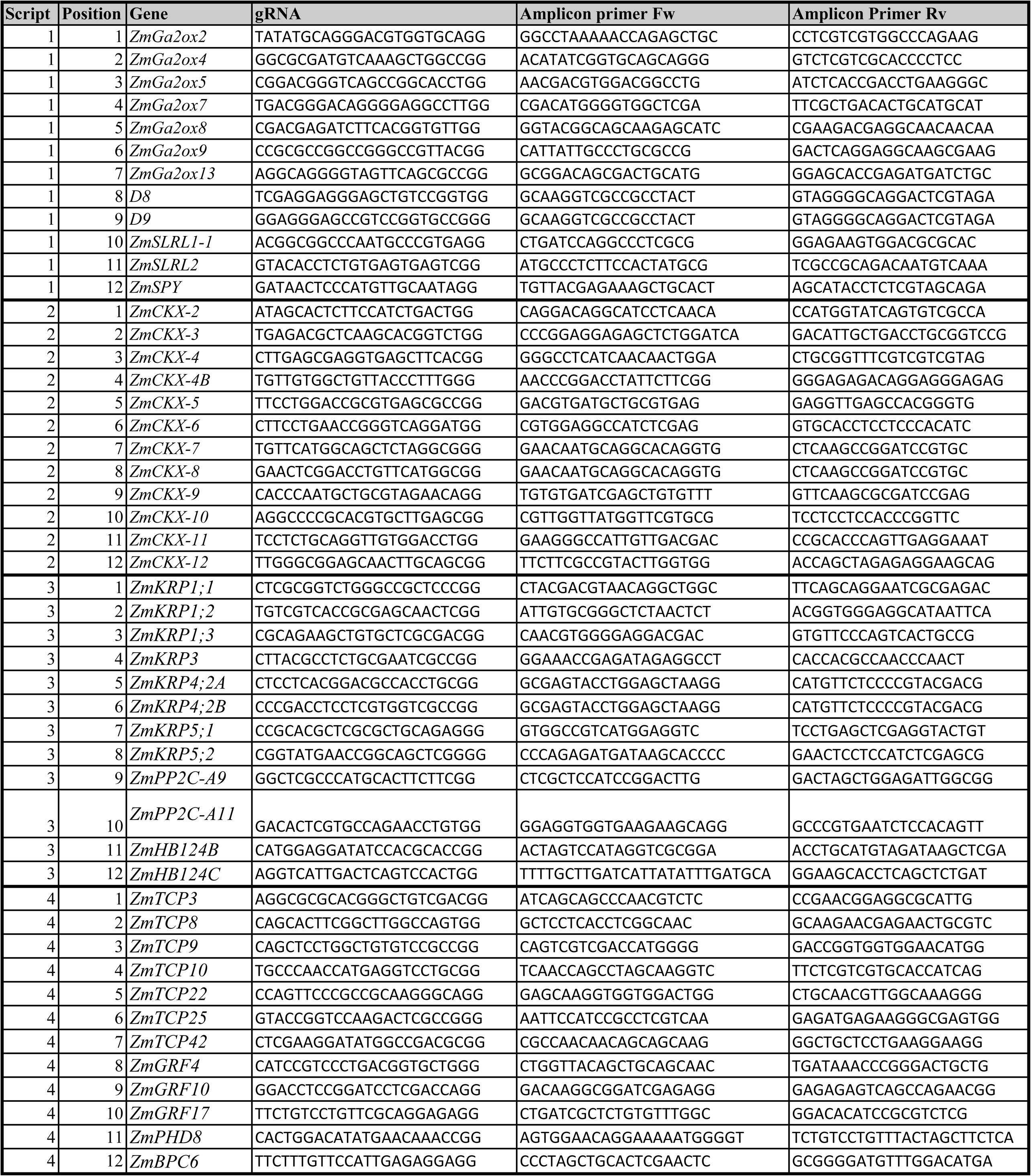
List of gRNAs and associated primer pairs for edit detection using HiPlex amplicon sequencing.

**Supplemental Table 3:** Plasmid overview.

| Full name | type | Cloning sites | Bacterial Selection | Plant Selection |
| --- | --- | --- | --- | --- |
| pGG-A-OsU3-Bbsl-ccdB-Bbsl-B | Entry vector | A-B | Ampicilin |  |
| pGG-B-OsU3-Bbsl-ccdB-Bbsl-C | Entry vector | B-C | Ampicilin |  |
| pGG-C-OsU3-Bbsl-ccdB-Bbsl-D | Entry vector | C-D | Ampicilin |  |
| pGG-D-OsU3-Bbsl-ccdB-Bbsl-E | Entry vector | D-E | Ampicilin |  |
| pGG-E-OsU3-Bbsl-ccdB-Bbsl-F | Entry vector | E-F | Ampicilin |  |
| pGG-F-OsU3-Bbsl-ccdB-Bbsl-G | Entry vector | F-G | Ampicilin |  |
| pGG-B-U1-OsU3-Bbsl-ccdB-Bbsl-C | Entry vector with U-linker | B-C | Ampicilin |  |
| pGG-C-U2-OsU3-Bbsl-ccdB-Bbsl-D | Entry vector with U-linker | C-D | Ampicilin |  |
| pGG-D-U3-OsU3-Bbsl-ccdB-Bbsl-E | Entry vector with U-linker | D-E | Ampicilin |  |
| pGG-E-U4-OsU3-Bbsl-ccdB-Bbsl-F | Entry vector with U-linker | E-F | Ampicilin |  |
| pGG-F-U5-OsU3-Bbsl-ccdB-Bbsl-G | Entry vector with U-linker | F-G | Ampicilin |  |
| pEN-2xTaU3 | PCR template |  | Kanamycin |  |
| pGG-A-Script1-entry1-B | Paired gRNA entry vector | A-B | Ampicilin |  |
| pGG-B-Script1-entry2-C | Paired gRNA entry vector | B-C | Ampicilin |  |
| pGG-C-Script1-entry3-D | Paired gRNA entry vector | C-D | Ampicilin |  |
| pGG-D-Script1-entry4-E | Paired gRNA entry vector | D-E | Ampicilin |  |
| pGG-E-Script1-entry5-F | Paired gRNA entry vector | E-F | Ampicilin |  |
| pGG-F-Script1-entry6-G | Paired gRNA entry vector | F-G | Ampicilin |  |
| pGG-A-Script2-entry1-B | Paired gRNA entry vector | A-B | Ampicilin |  |
| pGG-B-Script2-entry2-C | Paired gRNA entry vector | B-C | Ampicilin |  |
| pGG-C-Script2-entry3-D | Paired gRNA entry vector | C-D | Ampicilin |  |
| pGG-D-Script2-entry4-E | Paired gRNA entry vector | D-E | Ampicilin |  |
| pGG-E-Script2-entry5-F | Paired gRNA entry vector | E-F | Ampicilin |  |
| pGG-F-Script2-entry6-G | Paired gRNA entry vector | F-G | Ampicilin |  |
| pGG-A-Script2-entry1-B | Paired gRNA entry vector | A-B | Ampicilin |  |
| pGG-B-Script3-entry2-C | Paired gRNA entry vector | B-C | Ampicilin |  |
| pGG-C-Script3-entry3-D | Paired gRNA entry vector | C-D | Ampicilin |  |
| pGG-D-Script3-entry4-E | Paired gRNA entry vector | D-E | Ampicilin |  |
| pGG-E-Script3-entry5-F | Paired gRNA entry vector | E-F | Ampicilin |  |
| pGG-F-Script3-entry6-G | Paired gRNA entry vector | F-G | Ampicilin |  |
| pGG-A-Script4-entry1-B | Paired gRNA entry vector | A-B | Ampicilin |  |
| pGG-B-Script4-entry2-C | Paired gRNA entry vector | B-C | Ampicilin |  |
| pGG-C-Script4-entry3-D | Paired gRNA entry vector | C-D | Ampicilin |  |
| pGG-D-Script4-entry4-E | Paired gRNA entry vector | D-E | Ampicilin |  |
| pGG-E-Script4-entry5-F | Paired gRNA entry vector | E-F | Ampicilin |  |
| pGG-F-Script4-entry6-G | Paired gRNA entry vector | F-G | Ampicilin |  |
| pGG-A-Script5-entry1-B | Paired gRNA entry vector | A-B | Ampicilin |  |
| pGG-B-Script5-entry2-C | Paired gRNA entry vector | B-C | Ampicilin |  |
| pGG-C-Script5-entry3-D | Paired gRNA entry vector | C-D | Ampicilin |  |
| pGG-D-Script5-entry4-E | Paired gRNA entry vector | D-E | Ampicilin |  |
| pGG-E-Script5-entry5-F | Paired gRNA entry vector | E-F | Ampicilin |  |
| pGG-F-Script5-entry6-G | Paired gRNA entry vector | F-G | Ampicilin |  |
| pGGBb-AG | Destination vector | A-G | Spectinomycin | Bar |
| pGG-AG-Script1 | Expression vector | A-G | Spectinomycin | Bar |
| pGG-AG-Script2 | Expression vector | A-G | Spectinomycin | Bar |
| pGG-AG-Script3 | Expression vector | A-G | Spectinomycin | Bar |
| pGG-AG-Script4 | Expression vector | A-G | Spectinomycin | Bar |
| pGG-AG-Script5 | Expression vector | A-G | Spectinomycin | Bar |
| Supplemental Table 3: Plasmid overview. |  |  |  |  |

**Supplemental Table 4:**
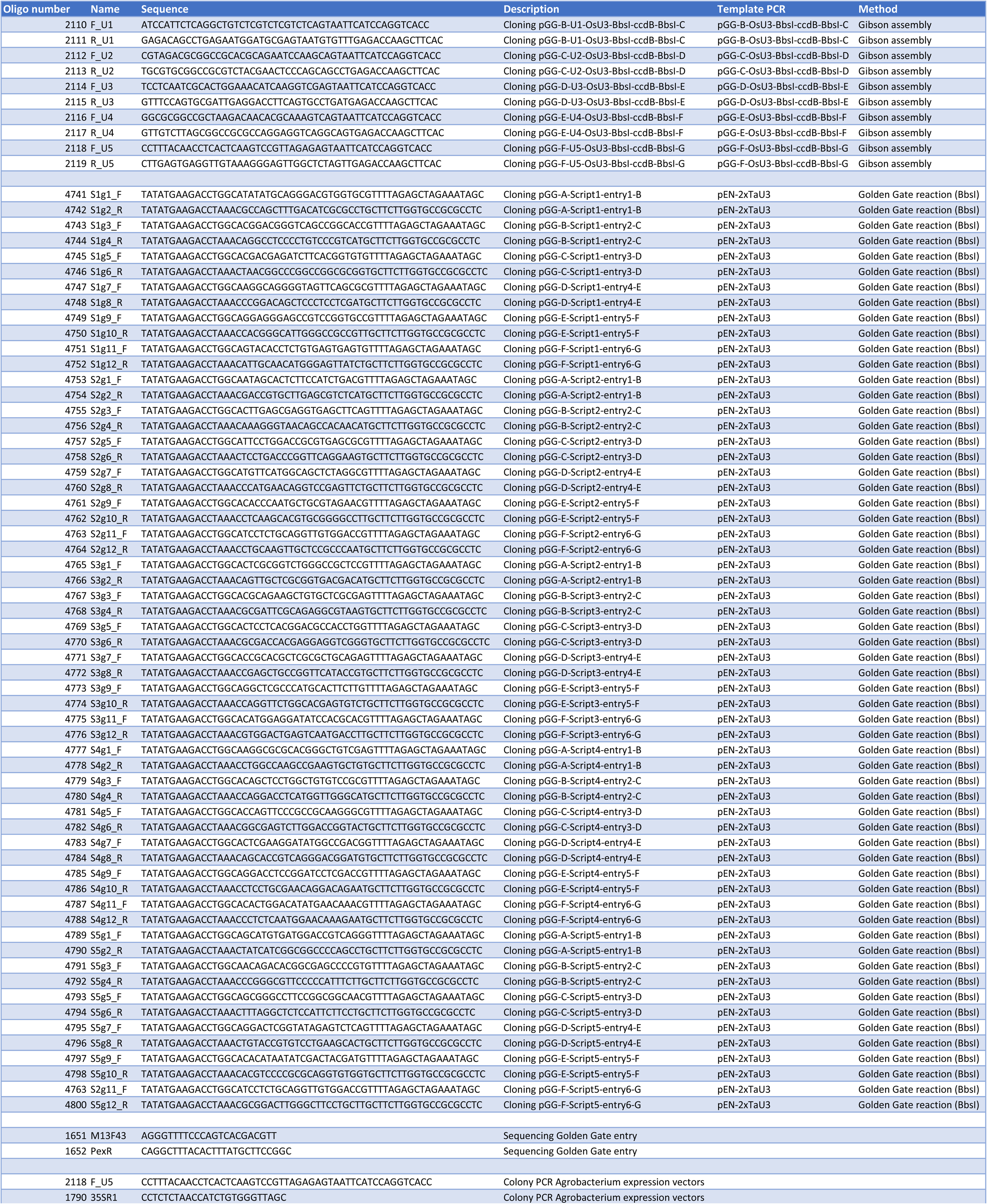
Primers used for plasmid building and sequencing.

## Acknowledgments

The authors would like to thank Lieven Sterck for helping in gene model curation, Mansour Karimi for helping with the cloning of EDITOR constructs, Pan Gong, Reinout Laureyns, and Ji Li for additional help with the selection of the genes.

